# Warming limits energetic investment in feeding and post-feeding metabolism in bumblebees

**DOI:** 10.64898/2026.06.11.731625

**Authors:** Natacha Rossi, Elizabeth Nicholls

## Abstract

Environmental warming is generally expected to increase metabolic demand in ectotherms. However, facultatively endothermic insects such as bumblebees regulate body temperature and may reduce thermogenic investment under warm conditions, potentially altering physiological performance and responses to climate change. We combined flow-through respirometry and infrared thermography to test how elevated ambient temperature (25 vs 35°C) affects feeding energetics and postprandial metabolism in the bumblebee *Bombus terrestris*. Bees maintained substantially lower thoracic temperature excess at 35°C than at 25°C, both before and during feeding. Feeding metabolic rate was also lower at 35°C and was strongly positively associated with thoracic temperature excess, indicating that feeding energetics were primarily explained by thermoregulatory state rather than ambient temperature alone. Elevated temperature reduced both the probability and energetic magnitude of specific dynamic action (SDA), including total SDA expenditure, early postprandial metabolism, and peak metabolic amplitude. In contrast, SDA duration and time to peak response showed little temperature dependence. Our results demonstrate that warming can suppress energetic expenditure in facultatively endothermic pollinators by limiting thermogenic investment and postprandial metabolic responses, potentially constraining the energetic flexibility underpinning foraging performance under climate warming.

## Introduction

Environmental warming is generally expected to elevate metabolic demand through temperature-dependent increases in physiological rates (Gillooly et al., 2001). However, facultatively endothermic insects such as bees actively regulate thoracic temperature through endogenous heat production in the flight muscles (Heinrich, 1979), meaning metabolic expenditure depends not only on ambient temperature, but also on thermogenic investment. Because bumblebees rely heavily on endogenous heat production to sustain foraging performance under cool conditions (Bishop & Armbruster, 1999), environmental warming may reduce rather than elevate energetic expenditure by suppressing thermogenesis as the risk of overheating rises.

In foraging bees, thoracic temperature strongly influences muscle performance, metabolic rate, and behavioural activity, even while bees are stationary and feeding at flowers (Coelho, 1991b, 1991a; Harrison & Fewell, 2002). Feeding therefore represents a distinct thermoregulatory context in which energetic expenditure reflects not only nutrient intake itself, but also the maintenance of elevated thoracic temperature during foraging. Feeding also initiates a sequence of post-ingestive processes, including crop emptying, proventricular transport, haemolymph sugar regulation, and nutrient assimilation, that collectively contribute to specific dynamic action (SDA): the postprandial elevation of metabolic rate above pre-feeding baseline (Blatt & Roces, 2001, 2002a, 2002b; McCue, 2006; Secor, 2009). In insects, SDA is best understood as a composite of transport, regulatory, and biosynthetic processes rather than a discrete digestive cost alone (McCue, 2006). In bees, these postprandial processes occur against a background of ongoing thermoregulation, making it difficult to disentangle digestion-associated metabolism from variation in thoracic temperature and baseline energetic state. Despite extensive work on insect thermoregulation and feeding energetics (Heinrich, 1974; Lee, 2019; Trier & Mattson, 2003), it remains unknown whether warming alters the probability, energetic magnitude, or temporal organisation of SDA responses in bees. Because SDA contributes to the overall energy budget of foraging insects, temperature-driven changes in SDA could alter resource requirements and energy acquisition, making it important for understanding pollinator responses to climate warming.

Bumblebees provide a well-suited system for addressing these questions. They are ecologically important pollinators, particularly in high latitude and altitude regions, and as a result, thought to be particularly vulnerable to climate warming (Soroye et al., 2020). During activity, bumblebees commonly maintain thoracic temperatures around 35–38°C, such that ambient temperatures approaching this range reduce thoracic temperature excess and the scope for further thermogenic heat production (Heinrich, 1979).

Previous work in honeybees and wasps has demonstrated that feeding metabolic rate can remain relatively independent of ambient temperature across moderate thermal conditions, consistent with active thermoregulatory compensation (Stabentheiner et al., 2012). However, comparable work in bumblebees remains sparse, and to our knowledge no insect study has combined measurements of feeding and postprandial metabolism with simultaneous thermal imaging of thermogenesis and behaviour. Bumblebees differ from honeybees in body size thermal ecology and reliance on endogenous heat production. Moreover, the pronounced intraspecific variation in bumblebee body size provides an opportunity to test whether ambient temperature modifies body mass dependence of feeding metabolic rate, a question relevant to growing evidence that warming influences body size in bees and other animals (Fitzgerald et al., 2022; Gérard et al., 2023; Theodorou et al., 2021; Gardner et al., 2011).

Here, we combine flow-through respirometry and infrared thermography to quantify metabolic rate, thoracic temperature, feeding behaviour, and postprandial metabolic responses under climate warming in the buff-tailed bumblebee, *Bombus terrestris*. *Bombus terrestris* is a managed bumblebee species available commercially for crop pollination, with a wide distribution across temperate Europe, the Mediterranean basin, the Near East, and parts of North Africa, including regions increasingly affected by extreme summer heat events (Beniston et al., 2017; C3S (Copernicus Climate Change Service), 2024; Rasmont et al., 2008; Widmer et al., 1998). Experimental and field evidence indicates that 20–25°C represent favourable foraging conditions, whereas temperatures in the low-to-mid 30s are more representative of elevated or heatwave-relevant conditions (Gérard et al., 2024; Kenna et al., 2021; Kwon & Saeed, 2003).

By integrating CO₂-based metabolic measurements, thoracic temperature dynamics, and behavioural observations across these two ecologically relevant conditions (25 vs 35°C), we treat feeding and post-feeding metabolism as a continuous thermoregulatory–metabolic process rather than as isolated physiological events, as in previous literature. We test three predictions (1) variation in feeding metabolic rate is explained primarily by thoracic temperature excess rather than ambient temperature, consistent with thermoregulatory mediation of energetic expenditure; (2) elevated temperature reduces both the probability and energetic magnitude of SDA responses, reflecting a contraction of thermogenic and postprandial metabolic; and (3) body-size dependence of metabolic rate is reduced under warming, as reduced thermogenic investment compresses physiological variation among individuals.

## Material and Methods

### Bees

We used three colonies of the buff-tailed bumblebee, *Bombus terrestris* (Koppert, UK). Each colony had *ad libitum* access to Koppert “nectar” solution within the colony and pollen was provided regularly directly to the nest. Colonies were individually connected to flight arenas measuring 41.5 x 60 x 36 cm that were illuminated using overhead LED panels (600 x 600mm, 3600 lm; Panel Value 600, Ledvance, RS Components, UK) and maintained on a 9h:15h light:dark cycle. Ambient temperature where the colonies were kept was maintained at approximately 25°C. Only active workers showing no visible signs of injury or abnormal behaviour were used in experiments.

### Temperature treatments

Experiments were conducted at either 25 °C or 35 °C in the respirometry room (separate from where the colonies were kept). Temperature treatments were assigned independently across bees within colonies. The order of temperature treatments across experimental days was alternated where possible to minimise temporal biases. Room temperature was maintained at either 25°C or 35°C (25.1 ± 0.2 °C or 35.1 ± 0.2 °C) using heaters and infrared lamps and the air pumped into the test chamber was monitored using a temperature probe sealed inside the chamber. Heating devices were positioned away from the chamber to avoid direct radiative heating of the bee during thermal recordings.

### Respirometry and thermal imaging setup

Metabolic rate and thoracic temperature were measured simultaneously using flow-through respirometry coupled with infrared thermography. Individual bees were placed in a custom chamber supplied with CO₂- and H₂O-scrubbed air (400 mL min⁻¹), while CO₂ production and dorsal thoracic temperature were recorded continuously throughout feeding. Full details of chamber construction, instrumentation and synchronisation procedures are provided in the Supplementary Methods.

### Feeding protocol

Individual bees underwent three feeding bouts within the respirometry chamber. Prior to each trial, bees were isolated for 1 h in the flight cage connected to the colony to reduce variability in feeding motivation and crop contents. The cage was illuminated from above using an LED panel, allowing bees to fly freely during this period, which promotes crop emptying through activity. Previous work shows substantial crop emptying occurs within the first hour after feeding, although complete emptying may require up to 2 h for concentrated solutions (Fournier et al., 2014). A 1 h deprivation period was therefore used to reduce crop contents while avoiding reduced activity observed during longer deprivation periods in pilot experiments.

Each feeding bout consisted of 10 µL of 35% (w/w) sucrose solution (Fisher, UK). The sucrose solution was delivered with a 10 µL Hamilton syringe (Hamilton, Reno, Nevada, USA) into a 3D-printed feeder (49 x 8 x 2.75 mm) positioned on the floor of the respirometry chamber with a magnet (3 x 1 mm), within the infrared window viewing aperture. The feeder was positioned at a fixed location within the infrared window viewing area to standardise thermal imaging during feeding. An identical feeder containing 10 µL of 35% (w/w) sucrose solution was placed in the control chamber for the entire duration of the trial to account for potential changes in CO_2_ levels due to the sucrose solution itself. The feeding volume falls within the range of nectar rewards encountered by insect pollinators (∼1–10 µL; S. Nicolson, 2007) while remaining below crop capacity and allowing precise control of intake. The sucrose concentration was selected to provide a high energetic reward while maintaining rapid ingestion rates, as higher concentrations increase viscosity and reduce feeding efficiency (Harder, 1986). The first feeding bout occurred 5 min after introduction of the bee into the test chamber to allow chamber gas equilibration and behavioural acclimation. The second and third feeding bouts occurred at 3 min intervals once CO_2_ concentrations returned to baseline. Feeders were visually inspected and weighed before and after each trial to confirm complete consumption. Bees that did not fully consume the sucrose solution were excluded from the dataset. Bees were also weighed before and after trial and subsequently euthanised by freezing at -20°C.

### Thermal imaging analysis

Thoracic surface temperature was quantified from infrared recordings collected during feeding trials. Background-corrected thoracic temperature excess was calculated as the difference between thoracic and background temperatures (ΔTthorax = Tthorax − Tbackground). Pre-feeding thoracic temperature was estimated immediately prior to feeding onset, and feeding temperature was estimated from measurements obtained during feeding. The feeding-associated thermal response was calculated as ΔΔT = ΔTfeeding − ΔTpre-feeding. Full details of thermal calibration, image processing and temperature extraction are provided in the Supplementary Methods.

### Respirometry data analysis

Metabolic rate (MR) was quantified from CO₂ production (V̇CO₂) measured using flow-through respirometry. For each feeding bout, pre-feeding MR was estimated immediately prior to feeding onset and feeding MR was estimated during the corresponding feeding interval after correcting for system washout delay. The feeding-associated metabolic response was calculated as ΔMR = MRfeeding − MRbaseline. Full details of synchronisation procedures, washout correction and respirometry calculations are provided in the Supplementary Methods.

### Specific dynamic action (SDA) quantification

Specific dynamic action (SDA) was quantified from post-feeding respirometry traces as the increase in metabolic rate associated with digestion and nutrient assimilation. For each feeding bout, post-feeding CO₂ production was compared with pre-feeding baseline metabolism to identify feeding-associated metabolic responses. Responses were classified as full when both a detectable post-feeding peak and subsequent recovery to baseline were observed, partial when recovery was not observed, and non-responses when no feeding-associated increase was detected.

SDA magnitude was quantified as excess CO₂ production above baseline, and additional metrics describing response dynamics included peak amplitude, time to peak, response duration and early-phase SDA (AUC₆₀). Excess CO₂ production was converted to energetic units assuming carbohydrate metabolism. Full details of SDA detection algorithms, threshold selection, energetic conversions and sensitivity analyses are provided in the Supplementary Methods.

### Replication statement

| Factor | Scale of inference | Scale at which factor was applied | Independent replicates |
| --- | --- | --- | --- |
| Ambient temperature (25°C vs 35°C) | Individual worker bees (*Bombus terrestris*) | Individual bee | 55 bees total across 3 colonies (25°C: 27 bees; 35°C: 28 bees) |
| Feeding bout (F1–F3) | Feeding responses within individuals | Repeated measurements within each bee | Up to 3 feeding bouts per bee; 136 feeding bouts analysed |
| Colony | Colony-level blocking factor (not primary inference scale) | Colony | 3 colonies |

### Statistical analysis

All analyses were conducted in R (v4.4.3) using mixed-effects modelling approaches. Because individual bees contributed repeated measurements across feeding bouts, BeeID was included as a random intercept in all repeated-measures models. Colony identity was included as a fixed blocking factor because only three colonies were studied. Ambient temperature (25°C versus 35°C), feeding bout (F1–F3), body mass, and feeding duration were included as explanatory variables where appropriate.

Feeding duration, thoracic temperature, and metabolic-rate responses were analysed using linear mixed-effects models. SDA response occurrence and the probability of obtaining a full SDA response were analysed using binomial generalised linear mixed models, whereas positive SDA metrics (total SDA energy, AUC₆₀, peak amplitude, and duration) were analysed using Gamma models with log links. Time-to-peak responses were analysed using log-transformed response times.

Models were initially fitted with biologically motivated interaction terms and simplified hierarchically while maintaining model structure. Final model formulations are provided in Table S1. Post hoc comparisons and estimated marginal means were obtained using emmeans with Holm-adjusted P-values where appropriate. Model assumptions were assessed using standard residual diagnostics. Additional details regarding variable transformations, model diagnostics, software packages and model-performance metrics are provided in the Supplementary Methods.

## Results

After excluding one observation with missing pre-trial body mass, the final dataset comprised 136 feeding bouts from 55 bees across three colonies. Mean pre-trial body mass was 0.187 ± 0.052 g (mean ± SD). Pre-trial body mass did not differ significantly among colonies or between temperature treatments, indicating that treatment-level differences in metabolic and thermal traits were not attributable to systematic body-size differences among groups (Supplementary Results). Repeated measures were obtained from most individuals: 36 bees contributed three feeding bouts, nine contributed two bouts, and ten contributed one bout. Sample sizes were well balanced across ambient temperatures and feeding bouts (25°C: F1 = 24, F2 = 25, F3 = 22; 35°C: F1 = 23, F2 = 21, F3 = 21).

### Feeding duration

Feeding duration was significantly affected by ambient temperature, feeding bout, body mass, and their three-way interaction (Figure 1; Temperature × Feeding bout × Body mass: χ²₂ = 6.55, p = .038). Feeding bouts were consistently shorter at 25°C (mean ± SD, F1= 23.0 ± 13.8 s; F2= 16.5 ± 8.3 s; F3= 15.1 ± 6.0 s) compared to 35°C (F1= 16.7 ± 7.4 s; F2= 13.9 ± 7.5 s; F3= 9.9 ± 3.0 s), within each feeding bout (F1: p = .004; F2: p = .033; F3: p = .002). At 25°C, F1 feeding bouts lasted longer than both F2 (p = .001) and F3 (p < .001), whereas F2 and F3 did not differ (p = .914). At 35°C, F1 lasted longer than F3 (p < .001), while other pairwise contrasts were not significant (all p ≥ .054).

**Figure 1.**
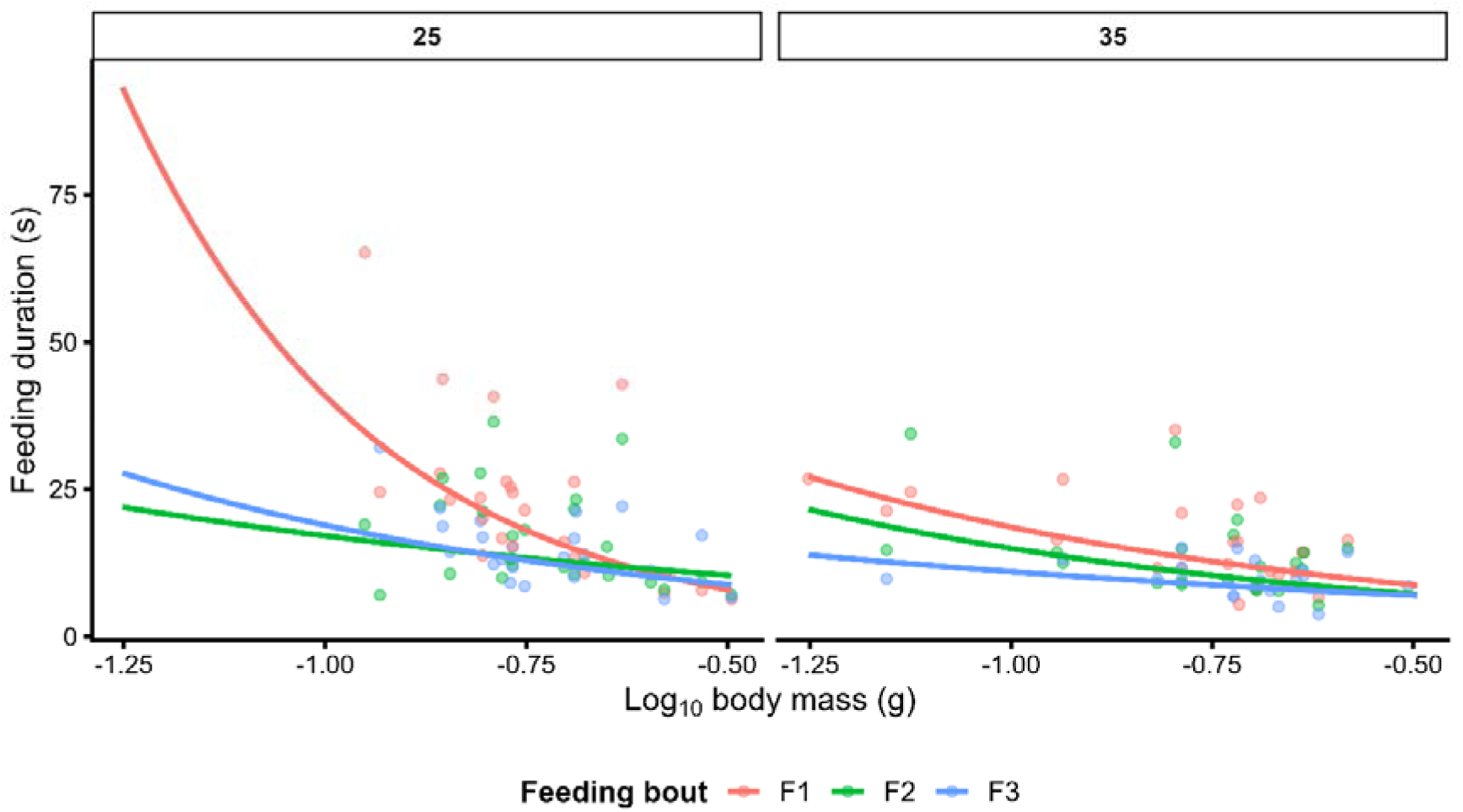
Predicted relationships between body mass and feeding duration across feeding bouts at 25°C and 35°C. Points represent raw observations and lines show model predictions from the final mixed-effects model.

Overall, larger bees completed feeding bouts more rapidly (χ²₁ = 19.50, p < .001), but the strength of this relationship depended on both feeding bout and ambient temperature (Figure 1). At 25°C, the negative relationship between body mass and feeding duration was stronger during F1 than F2 (p = .004) and marginally stronger than F3 (p = .050). By contrast, body-mass slopes did not differ among feeding bouts at 35°C (all p ≥ .611). The model explained a substantial proportion of variance in feeding duration (marginal R² = .428; conditional R² = .734), with moderate repeatability among individuals (ICC = .534).

### Thermal responses during feeding

Thermal responses during feeding was higher at 25°C than at 35°C (Figure 2A; χ²₁ = 52.08, p < .001; estimated contrast = 1.38 ± 0.19°C), ranging from 1.83 ± 1.31°C to 2.16 ± 1.04°C at 25°C and from 0.58 ± 0.30°C to 0.70 ± 0.39°C at 35°C. The relationship between body mass and thoracic temperature excess depended on ambient temperature (Figure 2B; Temperature × Body mass: χ²₁ = 6.98, p = .008), with larger bees exhibiting higher thoracic temperature excess at 25°C (3.75 ± 1.27, 95% CI = 1.20–6.31) but not at 35°C (−0.26 ± 0.82, 95% CI = −1.90 to 1.38).

**Figure 2.**
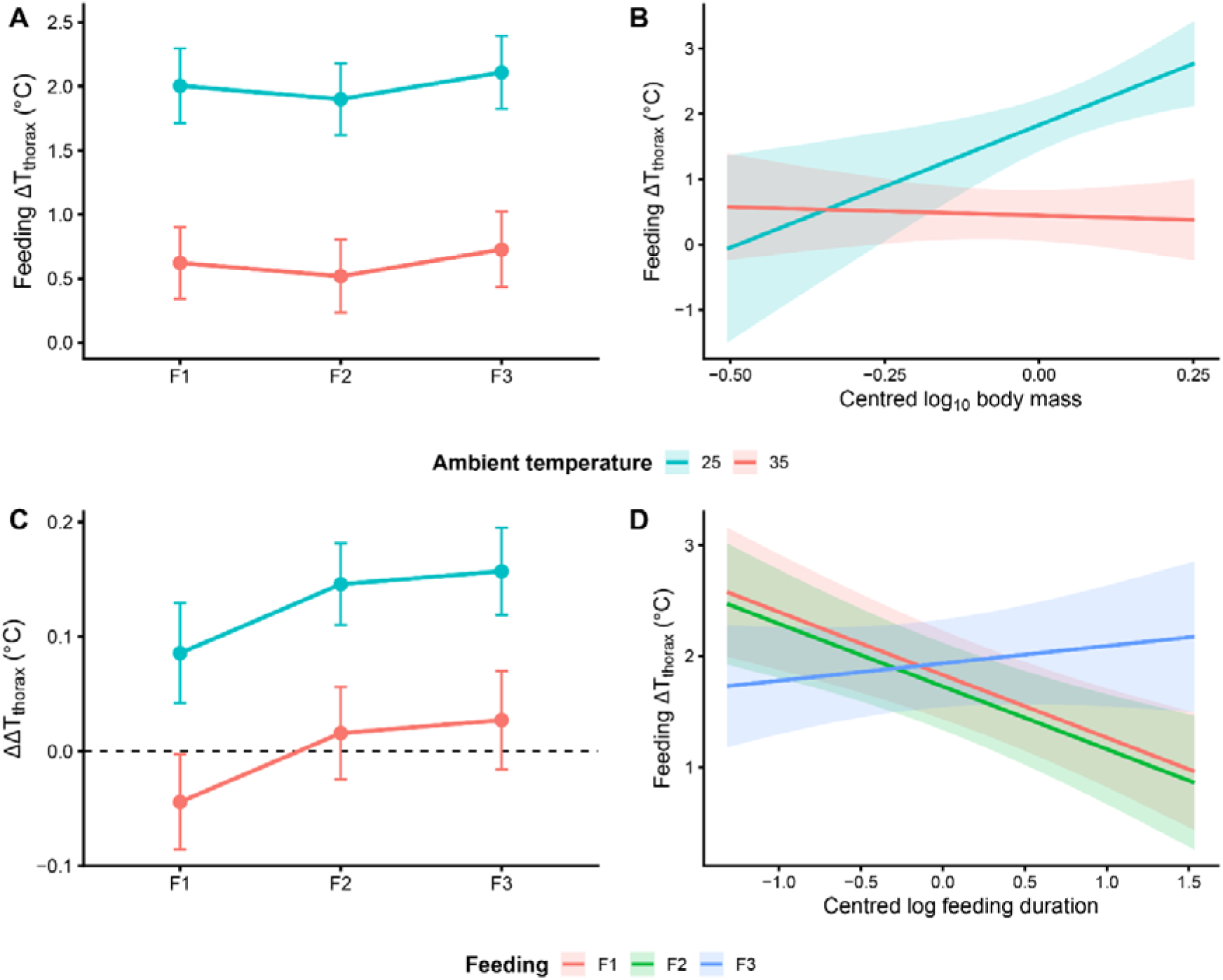
Feeding thoracic temperature responses and feeding-related interaction effects across ambient temperatures. (A) Feeding thoracic temperature excess (ΔT_thorax; thorax − background, °C) across feeding bouts (F1–F3) at 25°C and 35°C. Points represent estimated marginal means ± 95% confidence intervals from the final mixed-effects model. (B) Predicted relationship between feeding thoracic temperature excess and centred log₁₀ body mass across ambient temperatures. Lines represent model predictions and shaded ribbons represent 95% confidence intervals from the final mixed-effects model. (C) Change in thoracic temperature excess during feeding relative to pre-feeding levels (ΔΔT = ΔT_feed − ΔT_pre), with the dashed line indicating no change. Points represent estimated marginal means ± 95% confidence intervals from the final mixed-effects model. (D) Predicted relationship between feeding thoracic temperature excess and centred log feeding duration across feeding bouts. Lines represent model predictions and shaded ribbons represent 95% confidence intervals from the final mixed-effects model. Mixed-effects models included ambient temperature, feeding bout, body mass, feeding duration, and colony as fixed effects, with bee identity included as a random effect. Colours indicate ambient temperature in panels A–C and feeding bout in panel D.

Likewise, feeding duration interacted with feeding bout (Figure 2D; Feeding bout × Feeding duration: χ²₂ = 21.76, p < .001). Bees that fed for longer showed lower thoracic temperature excess during feeding in F1 (−0.57 ± 0.14, 95% CI = −0.84 to −0.29) and F2 (−0.56 ± 0.15, 95% CI = −0.86 to −0.27), whereas no clear relationship was detected during F3 (0.16 ± 0.17, 95% CI = −0.18 to 0.49). Feeding bout also affected thoracic temperature excess overall (χ²₂ = 7.75, p = .021), with lower values during F2 than F3 (post-hoc p = .019), whereas other feeding-bout contrasts were not significant (all p ≥ .38). The model explained substantial variation in feeding thoracic temperature excess (marginal R² = .541; conditional R² = .910), again with high repeatability among individuals (ICC = .803). Comparable patterns were observed for pre-feeding thoracic temperature excess (Supplementary Results; Figure S1).

Changes in thoracic temperature excess during feeding relative to pre-feeding were comparatively small. Mean Δthoracic temperature excess ranged from 0.09 ± 0.15°C to 0.15 ± 0.11°C at 25°C and from −0.002 ± 0.074°C to 0.051 ± 0.072°C at 35°C. Thermal change was greater at 25°C than at 35°C (Figure 2C; χ²₁ = 37.79, p < .001; estimated contrast = 0.13 ± 0.02°C) and also differed among feeding bouts (χ²₂ = 9.48, p = .009), with lower Δthermal responses during F1 than during F2 (post-hoc p = .033) and F3 (post-hoc p = .013), whereas F2 and F3 did not differ (p = .863). However, thermal change was also shaped by higher-order interactions involving body mass, feeding duration, ambient temperature and feeding bout, including Temperature × Body mass × Feeding duration (χ²₁ = 5.95, p = .015) and Feeding bout × Body mass × Feeding duration (χ²₂ = 15.64, p < .001). These interactions indicate that thermal change during feeding depended on the combined effects of body mass, feeding duration, ambient temperature, and feeding bout rather than showing a simple additive response. However, explanatory power was lower than for pre-feeding or feeding thoracic temperature excess (marginal R² = .370; conditional R² = .416), and repeatability among individuals was low (ICC = .073), indicating that feeding thermal change relative to pre-feeding was less consistently structured among individuals than absolute thoracic temperature excess.

### Metabolic rate before and during feeding

Feeding MR was substantially higher at 25°C than at 35°C across all feeding bouts (Figure 3A; χ²₁ = 63.22, p < .001), ranging from 4.04–4.71 mL h⁻¹ at 25°C and 1.22–1.56 mL h⁻¹ at 35°C. The highest feeding MR was observed during F3 at 25°C (4.71 ± 3.15 mL h⁻¹), whereas the lowest occurred during F3 at 35°C (1.22 ± 0.52 mL h⁻¹).

**Figure 3.**
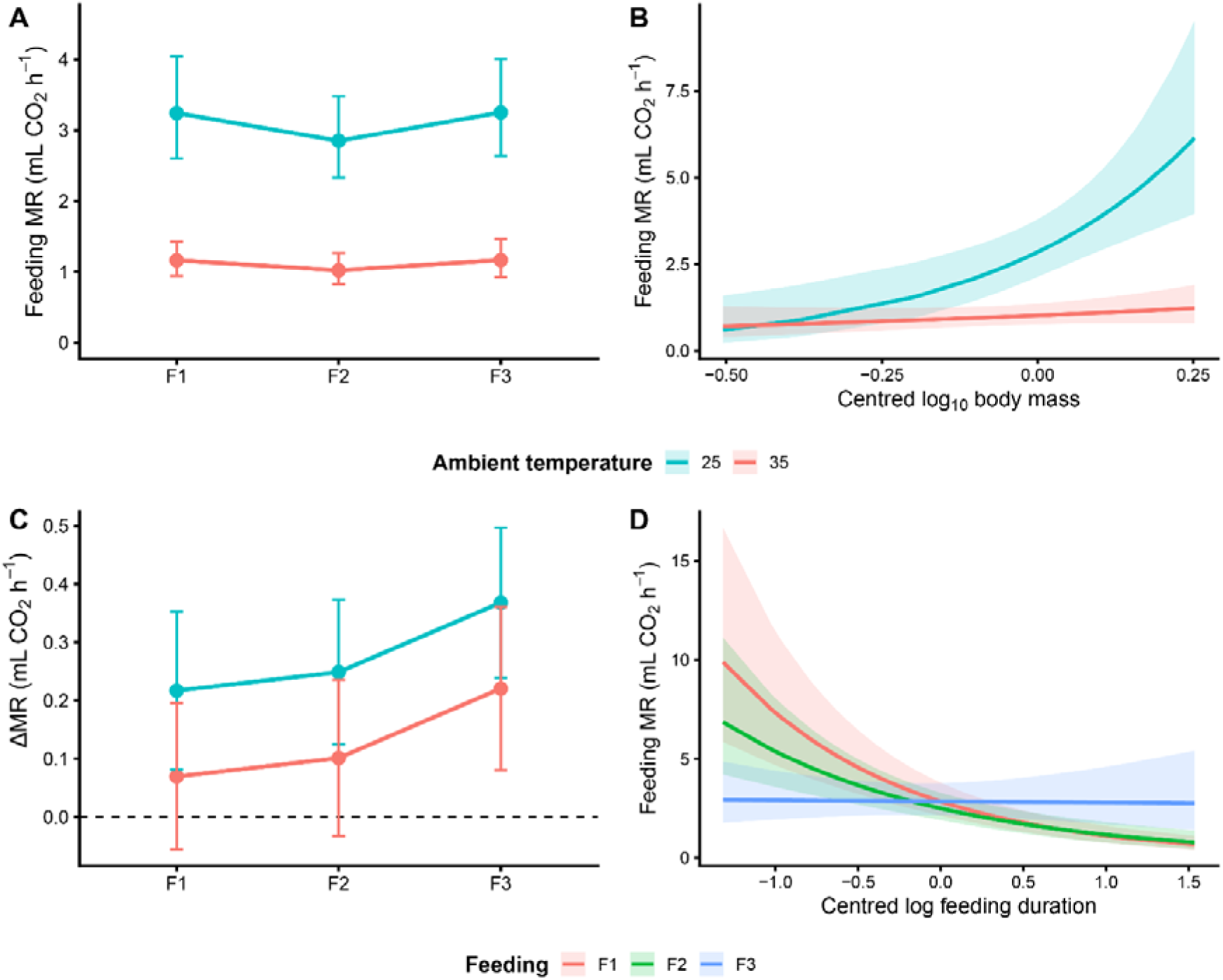
Feeding metabolic-rate responses and feeding-related interaction effects across ambient temperatures. (A) Feeding metabolic rate (MR; mL CO₂ h⁻¹) across feeding bouts (F1–F3) at 25°C and 35°C. Points represent estimated marginal means ± 95% confidence intervals from the final mixed-effects model. (B) Predicted relationship between feeding MR and centred log₁₀ body mass across ambient temperatures. Lines represent model predictions and shaded ribbons represent 95% confidence intervals from the final mixed-effects model. (C) Change in metabolic rate during feeding relative to pre-feeding levels (ΔMR = feeding − pre-feeding MR), with the dashed line indicating no change. Points represent estimated marginal means ± 95% confidence intervals from the final mixed-effects model. (D) Predicted relationship between feeding MR and centred log feeding duration across feeding bouts. Lines represent model predictions and shaded ribbons represent 95% confidence intervals from the final mixed-effects model. Mixed-effects models included ambient temperature, feeding bout, body mass, feeding duration, and colony as fixed effects, with bee identity included as a random effect. Colours indicate ambient temperature in panels A–C and feeding bout in panel D.

Feeding MR increased with body mass overall (χ²₁ = 13.38, p < .001), although this relationship depended on ambient temperature (Figure 3B; Temperature × Body mass: χ²₁ = 5.38, p = .020). Feeding MR increased strongly with body mass at 25°C (1.32 ± 0.37, 95% CI = 0.59–2.06), whereas the relationship was weaker and non-significant at 35°C (0.32 ± 0.25, 95% CI = −0.19 to 0.82).

Feeding MR declined with increasing feeding duration (χ²₁ = 22.23, p < .001), but this effect differed among feeding bouts (Figure 3D; Feeding bout × Feeding duration: χ²₂ = 22.45, p < .001). Feeding duration was negatively associated with feeding MR during F1 (−0.41 ± 0.07, p < .001) and F2 (−0.33 ± 0.07, p < .001), but not during F3 (−0.01 ± 0.08, p = .91). The negative association between feeding duration and feeding MR also weakened as body mass increased (Body mass × Feeding duration: χ²₁ = 6.48, p = .011). The model explained substantial variation in feeding MR (marginal R² = .620; conditional R² = .798), with moderate repeatability among individuals (ICC = .467).

The increase in MR during feeding relative to pre-feeding levels (ΔMR) was modest, ranging from 0.20–0.43 mL h⁻¹ at 25°C and 0.09–0.18 mL h⁻¹ at 35°C. ΔMR was significantly higher at 25°C than at 35°C (Figure 3C; χ²₁ = 4.12, p = .042), whereas feeding bout (χ²₂ = 4.24, p = .120), body mass and feeding duration were not significant predictors (both p ≥ .15). Explanatory power (marginal R² = .127; conditional R² = .215), and repeatability among individuals (ICC = .101) were substantially lower than for absolute feeding MR.

Comparable patterns were observed for pre-feeding MR, and all sensitivity analyses (body-mass filtering, alternative scaling approaches, Q₁₀ correction and mass-specific MR analyses) supported the primary conclusion that metabolic rates were consistently higher at 25°C than at 35°C (Supplementary Results; Figures S2–S3, Tables S2–S3).

### Effect of thoracic temperature excess on metabolic rate

Feeding MR increased strongly with thoracic temperature excess during feeding (Figure 4; χ²₁ = 175.84, p < .001) and the strength of this increase depended on ambient temperature (dT × Temperature: χ²₁ = 10.43, p = .001). There was a steeper positive association at 35°C (0.465 ± 0.054, 95% CI = 0.358–0.571) than at 25°C (0.297 ± 0.015, 95% CI = 0.266–0.327; slope contrast: p = .002).

**Figure 4.**
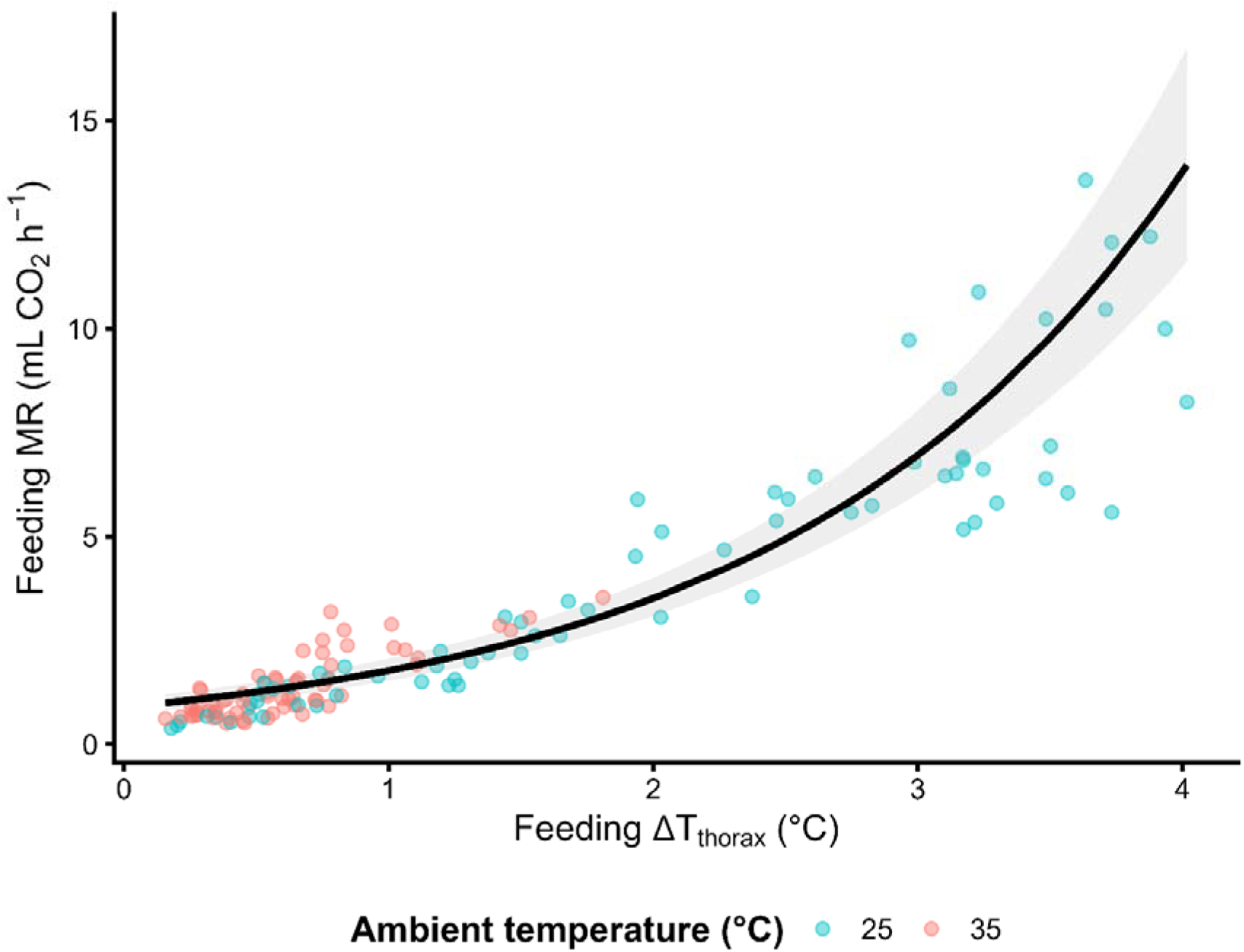
Relationship between thoracic temperature excess and feeding metabolic rate. Feeding metabolic rate (MR; mL CO₂ h⁻¹) increased strongly with thoracic temperature excess during feeding (ΔTthorax = thorax − background, °C). Points represent individual feeding observations coloured by ambient temperature treatment (25°C vs 35°C). The solid black line shows the fitted relationship predicted from the final mixed-effects mechanistic model, with shaded ribbons indicating 95% confidence intervals. Predictions are shown on the original MR scale following back-transformation from log₁₀-transformed model estimates. The model included ambient temperature, feeding bout, body mass, feeding duration, colony, and retained interaction terms involving thoracic temperature excess, with bee identity included as a random effect.

The relationship between thoracic temperature excess and feeding MR also depended on feeding duration (dT × Feeding duration: χ²₁ = 5.52, p = .019), tending to strengthen during longer feeding bouts, although pairwise contrasts among representative feeding durations were marginal after adjustment (Tukey-adjusted p = .059).

Once thoracic temperature excess was included in a model of feeding MR, feeding bout no longer explained feeding MR (χ²₂ = 1.04, p = .594), suggesting that feeding-bout differences in metabolic rate were largely accounted for by thoracic thermal state itself. The final model explained most variation in feeding MR (marginal R² = .903; conditional R² = .909), while residual repeatability among individuals was low (ICC = .061).

### Magnitude of specific dynamic action responses to feeding

Specific dynamic action (SDA) responses, defined as post-feeding increases in metabolic rate, were strongly influenced by ambient temperature. Across 137 feeding bouts, detectable SDA responses occurred in 109 cases, with subsequent analyses conducted on variable-specific subsets depending on data completeness and response characteristics.

The probability of detecting an SDA response was significantly higher at 25°C than at 35°C (Figure 5A; χ²₁ = 6.59, p = .010). Response occurrence remained consistently high across feeding bouts at 25°C (87.5%-90.9%) but was lower and more variable at 35°C (60.9%-81.8%). Feeding bout, body mass, feeding duration, and colony identity did not significantly affect response occurrence (all p ≥ .22).

**Figure 5.**
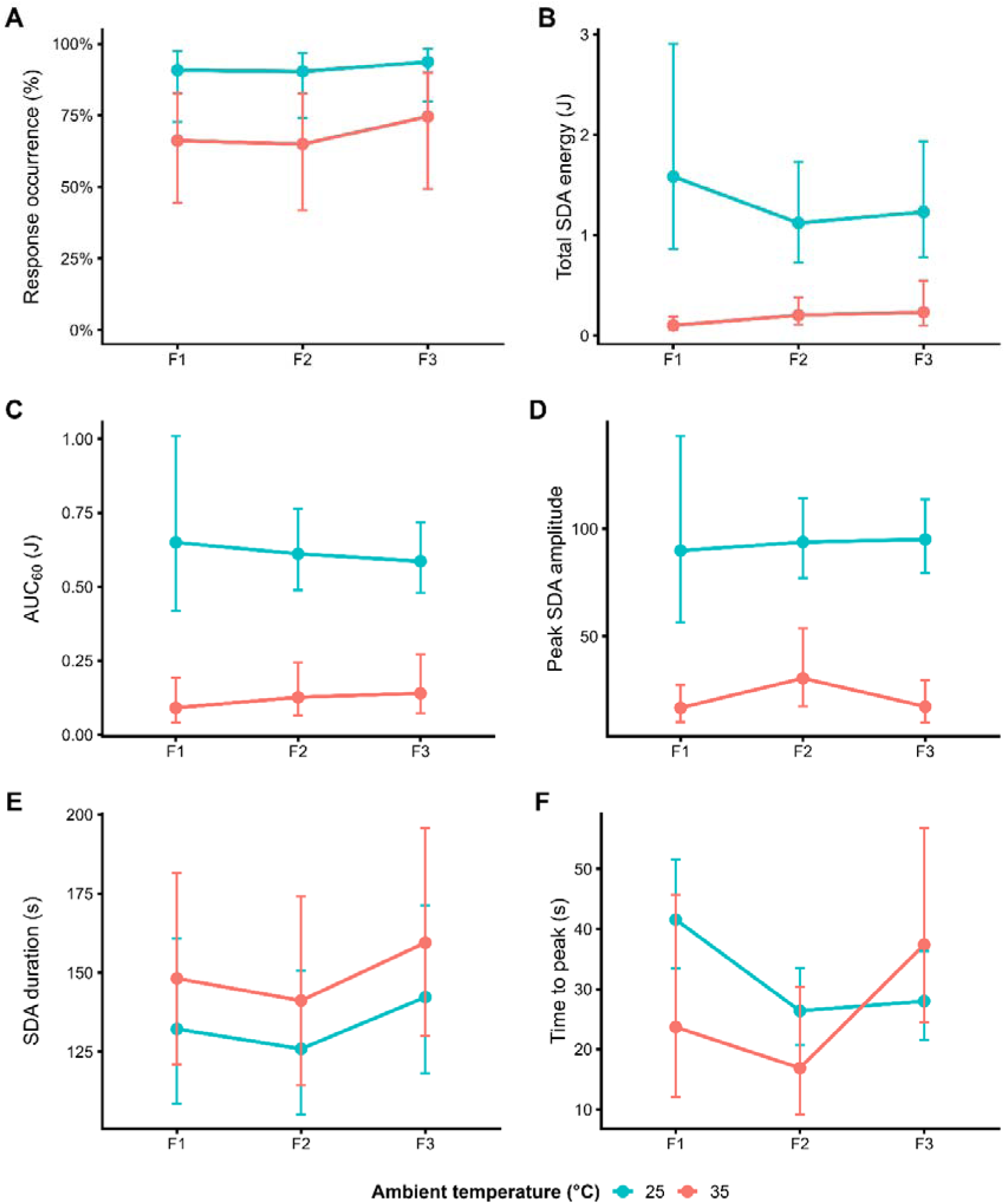
Specific dynamic action (SDA) responses following feeding at 25°C and 35°C across three sequential feeding bouts (F1–F3). (A) Probability of detecting an SDA response (%). (B) Total SDA energetic expenditure (strict SDA scope energy, J). (C) Early-phase SDA magnitude quantified as the area under the post-feeding metabolic response curve during the first 60 s following feeding (AUC₆₀, J). (D) Peak SDA amplitude. (E) SDA duration (s). (F) Time required to reach peak metabolic response following feeding (s). Points represent estimated marginal means ± 95% confidence intervals from final mixed-effects models including ambient temperature, feeding bout, body mass, feeding duration, and colony as fixed effects, with bee identity as a random effect. Colours indicate ambient temperature (25°C vs 35°C).

Ambient temperature also strongly affected the energetic magnitude of SDA responses. Among feeding bouts producing quantifiable SDA responses (n = 107), total SDA energetic expenditure (strict SDA scope energy) was substantially lower at 35°C than at 25°C (Figure 5B; χ²₁ = 49.86, p < .001). Median SDA energy ranged from 1.03–1.15 J at 25°C but only 0.05–0.15 J at 35°C. Feeding bout itself did not significantly affect SDA scope (χ²₂ = 0.98, p = .614), although the final selected model retained significant higher-order interactions involving ambient temperature, feeding bout, body mass, and feeding duration (Table S1). Post hoc analyses indicated that feeding duration was negatively associated with SDA scope during F1 at 25°C (−1.30 ± 0.64, 95% CI = −2.56 to −0.05), whereas no other feeding-duration trends were clearly supported.

Early-phase SDA magnitude, quantified as the area under the post-feeding metabolic response curve during the first 60 s following feeding (AUC₆₀; n = 127), showed an even stronger temperature dependence (Figure 5C; χ²₁ = 55.78, p < .001). Median AUC₆₀ values ranged from 0.57–0.67 J at 25°C but only 0.01–0.07 J at 35°C. The final selected model retained significant log Body mass × Feeding duration (χ²₁ = 7.30, p = .007) and Temperature × Feeding bout × Body mass interactions (χ²₂ = 8.45, p = .015), indicating that early SDA responses depended on the combined effects of body size, feeding behaviour, and thermal environment. Post hoc analyses indicated that the increase in AUC₆₀ with body mass was strongest during F3 at 25°C, whereas no corresponding body-mass associated increase was evident at 35°C (p = .0045).

Peak SDA amplitude (maximum post-feeding metabolic elevation; n = 109) showed a similar pattern, with substantially larger post-feeding metabolic elevations at 25°C than 35°C (Figure 5D; χ²₁ = 55.86, p < .001). Median peak amplitudes ranged from 90.5–108.2 at 25°C but only 14.8–23.0 at 35°C. Peak amplitude also varied significantly with body mass overall (χ²₁ = 6.83, p = .009), however the final selected model retained several higher-order interactions involving feeding bout, feeding duration, body mass, and ambient temperature. Post hoc analyses indicated that the negative relationship between feeding duration and SDA peak amplitude at 35°C was strongest in smaller individuals and weakened with increasing body mass. By contrast, feeding duration showed weakly positive or near-zero relationships with peak amplitude at 25°C. In addition, positive body-mass scaling of peak amplitude was strongest during F3 relative to F2 (p < .001).

### Timing of SDA responses to feeding

In contrast to SDA magnitude, the temporal characteristics of SDA responses did not vary with ambient temperature (Figure 5E; χ²₁ = 1.20, p = .273), with median SDA durations ranging from 121–140 s at 25°C and 121–178 s at 35°C. SDA duration (n = 108 recovered responses) was also unaffected by feeding bout (χ²₂ = 1.08, p = .583), body mass (χ²₁ = 0.13, p = .723) or feeding duration (χ²₁ = 0.004, p = .947).

Similarly, time to peak metabolic response showed weak and less consistent treatment dependence (Figure 5F). Although neither ambient temperature (χ²₁ = 1.41, p = .236) nor feeding bout (χ²₂ = 5.67, p = .059) significantly affected time-to-peak overall, the final selected model retained interaction terms involving feeding bout, body mass, and feeding duration. However, post hoc analyses did not reveal clear pairwise differences among feeding bouts. Median time-to-peak values ranged from 29–37 s at 25°C and 11–21 s at 35°C.

## Discussion

Predicting how environmental warming affects insect energetics is central to understanding the ecological consequences of climate change. In ectothermic animals, warming is generally expected to increase metabolic rate through passive thermal effects on biochemical and physiological processes (Clarke, 2004; Gillooly et al., 2001). However, facultatively endothermic insects such as bumblebees regulate thoracic temperature through active heat production, meaning that warming may instead reduce thermogenic investment and alter the energetic scope available for behaviour (Glass & Harrison, 2024; Heinrich, 1979). Despite extensive work on flight energetics and thermoregulation, much less is known about how warming affects the energetics of feeding and postprandial metabolism during flower visitation. By combining flow-through respirometry with infrared thermography, here we show that elevated ambient temperature constrained rather than accelerated energetic investment in feeding and postprandial metabolism in the bumblebee *Bombus terrestris*. Relative to bees at 25°C, at heat-wave relevant temperatures of 35°C, bees maintained lower thoracic temperature excess, exhibited lower feeding metabolic rates, and showed substantially reduced specific dynamic action (SDA) responses indicating reduced energetic investment in post-feeding metabolism.

The most striking result was the suppression of SDA at 35°C. Direct evidence that elevated temperatures attenuate postprandial metabolism in insects remains scarce, and we found no comparable studies examining temperature-dependent SDA responses in Hymenoptera. The closest insect comparison is the blood-feeding tsetse fly *Glossina brevipalpis*, in which warming reduced SDA magnitude but also substantially shortened digestion duration (McCue et al., 2016). In contrast, warming in *B. terrestris* primarily reduced the probability and energetic magnitude of in SDA responses, while leaving temporal structure largely unchanged.

SDA represents the postprandial elevation in metabolism associated with nutrient handling and processing (Secor, 2009). Although SDA accounted for only a modest proportion of meal energy, detectable responses were substantially less frequent at 35°C and, when present, 10-fold lower than at 25°C. Each 10 µL sucrose meal contained approximately 66 J of chemical energy, placing SDA at roughly 1–2% of meal energy at 25°C but only ∼0.1–0.3% at 35°C. In worker bees, nectar may also be temporarily retained within the crop prior to downstream processing, meaning that SDA likely reflects a composite of transport, regulatory, storage, and assimilation-related processes rather than digestive assimilation alone (Blatt & Roces, 2002a, 2002b; Fournier et al., 2014). These SDA values are low relative to available comparative data, including blood-feeding tsetse flies (16.7% at 25°C vs 4.6% at 35°C) (McCue et al., 2016) and most vertebrate ectotherms (Secor, 2009) where SDA typically accounts for 10-30% of meal energy. However, direct comparisons should be interpreted cautiously because SDA magnitude depends strongly on meal size, composition, temperature, and methodological definitions (Goodrich et al., 2024; McCue, 2006; Secor, 2009).

The stability of SDA timing, despite large reductions in response magnitude suggests that elevated temperature did not fundamentally reorganise postprandial processing. Instead, bees appeared to invest less energy in those processes during the period studied. One possibility is that warming reduced the intensity of immediate post-ingestive nutrient processing, such as crop emptying or sugar transport, thereby lowering SDA and immediate availability of carbohydrates post-feeding, without necessarily impacting subsequent nutrient assimilation (Blatt & Roces, 2001, 2002a, 2002b; Fournier et al., 2014).

Alternatively, or in addition, elevated temperatures may have reduced the energetic capacity available for postprandial processing. In a variety of ectotherms, warming can reduce the residual aerobic scope available after feeding even when digestive processing continues successfully (Bihun et al., 2024; Frisk et al., 2013). Under this scenario, reduced SDA would likely reflect a strategic or physiological reduction in energetic investment rather than impaired digestion *per se*. This interpretation is consistent with the broader thermoregulatory patterns observed in our study, where bees at 35°C exhibited lower thoracic temperature excess and lower feeding metabolic rates. If bees approaching their upper thermal limits have reduced capacity to tolerate additional metabolically generated heat, postprandial processes may be downregulated to minimise further thermal loading. Because we did not directly measure crop-emptying dynamics, haemolymph sugar regulation, assimilation efficiency, or aerobic capacity, these interpretations remain provisional. Relative humidity was also not independently regulated after air scrubbing in our set up, and there is a possibility that dehydration may also interact with thermal stress.

Thoracic temperature excess was a stronger predictor of feeding metabolic rate than ambient temperature, consistent with evidence that thoracic temperature determines the energetic and mechanical capacity of bees during activity (Coelho, 1991b; Harrison & Fewell, 2002; Kenna et al., 2021; Woodrow et al., 2025). Our results extend this relationship to feeding, a behavioural context in which bees are stationary rather than flying. Even during nectar consumption, metabolic expenditure remained tightly linked to thermoregulatory state, suggesting that bees maintain a level of thoracic activation while exploiting floral resources, likely preserving readiness for take-off, flower-to-flower movement, or rapid behavioural responses. This pattern is consistent with honeybee studies showing that energetic expenditure during feeding is governed primarily by thermoregulatory state and can be adjusted according to reward and environmental conditions rather than ambient temperature alone (Moffatt, 2001; Stabentheiner et al., 2012; Stabentheiner & Kovac, 2014, 2016a). The thermoregulatory dependence persisted under heatwave-relevant temperatures, but with a clear reduction in thermogenic and metabolic capacity at 35°C.

Bumblebees typically maintain thoracic temperatures within a narrow range that optimises flight-muscle performance (Heinrich, 1979), yet as ambient temperatures approach these operating temperatures, the scope for further thermogenesis declines. Elevated temperatures increase the risk of overheating and proximity to critical thermal limits (Sepulveda & Goulson, 2023), which can lead to protein denaturation, cell death, loss of cognitive function (Sepúlveda-Rodríguez et al., 2024) and loss of muscle co-ordination (Woodrow et al., 2025). Consequently, the lower metabolic rates observed at 35°C likely reflect active reduction of thermogenesis rather than a simple reduction in physiological demand. Unlike classical ectotherms, where warming passively accelerates metabolic processes (Clarke, 2004; Gillooly et al., 2001), much of bumblebee metabolic expenditure reflects regulated thermogenesis. As ambient temperature approaches thoracic operating temperatures, bees appear to reduce endogenous heat production, producing lower rather than higher whole-animal metabolic rates. Warming may therefore constrain performance in endothermic insects not because metabolism becomes excessively high, but because the physiological scope for regulated heat production becomes compressed.

Feeding behaviour changed in parallel with these physiological responses. Feeding bouts were consistently shorter at 35°C, and bees with higher metabolic rates tended to feed more quickly, particularly during early bouts. Shorter feeding at elevated temperatures could reflect altered motivation, reduced tolerance of warm stationary conditions, or changes in ingestion dynamics. Regardless of mechanisms, our results demonstrate that nectar feeding portion of a floral visit itself is thermally sensitive. Because nectar handling time influences foraging efficiency and pollination dynamics (Harder, 1986; Spaethe & Weidenmüller, 2002), temperature-driven shifts in feeding duration may scale up to affect flower visitation patterns and resource intake.

Patterns associated with body mass further support a thermoregulatory interpretation, although they should be treated cautiously. Larger bees showed higher thoracic temperature excess and higher feeding metabolic rates at 25°C, but these relationships were weaker at 35°C. This is consistent with larger bees having greater thermogenic capacity under cooler conditions (Bishop & Armbruster, 1999; Heinrich, 1979), while elevated temperatures compress physiological variation as bees approach a common thermal ceiling. However, supplementary analyses showed that the apparent temperature dependence of metabolic scaling was partly influenced by high-mass individuals.

The reduction in feeding and postprandial energetic investment at elevated ambient temperatures has important ecological implications. If warming reduces feeding duration, thermogenic investment, and post-feeding metabolic processing, it may reduce the energetic profitability of floral visits and ultimately affect nectar intake, resource transport and colony provisioning. These thermoregulatory constraints are also likely to affect energetically demanding behaviours not measured here, including flight and floral buzzing. Buzz pollination relies on rapid activation of the flight muscles to generate high-frequency vibrations and imposes substantial energetic costs on bees (De Luca & Vallejo-Marín, 2013; Rossi et al., 2026), while recent evidence that vibration performance declines at higher temperatures (Woodrow et al., 2025) is consistent with the contraction of thermogenic scope observed here. Together, these findings suggest that warming may reduce the energetic and mechanical capacity required to exploit floral resources with high behavioural demands.

Our experimental design necessarily constrained sustained flight and behavioural thermoregulation, and did not capture natural variation in solar radiation, wind, floral morphology, or resource distribution. In the field, bees may mitigate thermal stress through behavioural adjustments such as altered foraging schedules, shade use, evaporative cooling, or flower choice. Future studies should therefore examine how these behavioural responses interact with thermoregulatory and postprandial energetics under natural conditions.

More broadly, our findings suggest that facultatively endothermic pollinators may experience thermal vulnerability differently from strictly ectothermic insects. Rather than exhibiting simple metabolic acceleration under warming, bumblebees appear to experience a contraction of thermogenic and energetic scope for behaviours associated with foraging.

As heatwave-relevant temperatures become increasingly common across Europe (Beniston et al., 2017; C3S (Copernicus Climate Change Service), 2024; Intergovernmental Panel on Climate Change (IPCC), 2023), digestive and post-ingestive energetic processes should be considered alongside flight, cognition and foraging behaviour when predicting pollinator responses to climate change. By combining flow-through respirometry, infrared thermography, and behavioural measurements, we were able to examine feeding and postprandial metabolism as a continuous thermoregulatory process rather than as isolated physiological phases. Elevated ambient temperature reduced thoracic temperature excess, feeding metabolic rate, and the probability and magnitude of SDA responses in *B. terrestris*, revealing that warming constrains energetic investment in both nectar feeding and postprandial metabolism. Predicting pollinator resilience under climate warming will therefore require consideration not only of ambient temperature, but also for the thermoregulatory limits that shape energetic scope in facultatively endothermic bees.

## Acknowledgements

We thank Charlie Woodrow for sharing his emissivity measurements of *B. terrestris* thoraxes that we used in our study. We also thank Solomon Ngotho for helping with building the microcontroller used to synchronise the thermal camera and respirometer. The first author used ChatGPT (OpenAI) to assist with code development, troubleshooting, data analysis workflows, and editing of manuscript text. All code, analyses, interpretations, and final manuscript content were reviewed, verified, and approved by the authors, who take full responsibility for the accuracy and integrity of the work.

## Contributions

NR and EN conceived the ideas and designed methodology; NR collected the data; NR analysed the data; NR led the writing of the manuscript. All authors contributed critically to the drafts and gave final approval for publication.

## Funding Information

This study was supported by a UKRI Future Leaders Fellowship (MR/T021691/1) and a Royal Society Research Grant (RGS\R2\242536) awarded to EN.

## Data availability statement

Data available from Zenodo: https://doi.org/10.5281/zenodo.20559308 (Rossi, 2026).

## Notes

https://doi.org/10.5281/zenodo.20593389

### Competing Interest Statement

The authors have declared no competing interest.

