## Supplementary Material for "Warming limits energetic investment in feeding and post-feeding metabolism in bumblebees"

Supplementary Methods:

*Respirometry and thermal imaging setup*

The respirometry chamber was a custom-built Perspex cylinder (10 cm length x 5 cm diameter) sealed at each end using 3D-printed plugs secured by rubber bands. An extra 3D-printed funnel was sealed on the chamber wall to mount an infrared inspection window (viewing aperture diameter: 45 mm; FLIR, RS Components, UK) allowing thermal recording of the bee with an infrared camera (FLIR A700 camera with a FLIR lens f = 18 mm (24°) f/1.0). Air scrubbed of CO_2_ and H_2_O was pumped through the chamber at a consistent rate of 400 ml/min regulated by mass flow controller (GFC17; Aalborg, NY, USA) positioned immediately prior to the test chamber. A second identical mass flow controller was positioned immediately prior to a parallel control chamber identical to the test chamber. Relative humidity inside the chamber was not independently regulated following air scrubbing. Air exiting the test and control chambers was analysed simultaneously for CO_2_ concentrations using an infrared LI-7000 CO_2_/H_2_O analyser (LI-COR Biosciences, Lincoln, Nebraska, USA).

The thermal camera was positioned perpendicular to the infrared window at a fixed distance of 18 cm from the chamber, providing a dorsal view of the bee during feeding. The feeder position ensured the bee remained within the thermal camera field of view while feeding.

Bees were placed individually inside the test chamber. They were able to walk freely within the chamber but could not sustain flight. The chamber floor was lined with disposable aluminium foil to minimise contamination by residual sucrose between trials.

Prior to each trial, the respirometry system and thermal camera were synchronised using a manual visual cue. A microcontroller-triggered synchronisation pulse simultaneously activated an audible buzzer and sent a voltage signal to the gas analysis system.

*Thermal imaging analysis*

As described above, thermal recordings were obtained using a top-mounted infrared camera focused on the feeder region of the respirometry chamber. Thoracic surface temperature was quantified from dorsal thorax measurements obtained while bees were visible within the thermal camera field of view. Only periods in which the thorax was clearly visible and unobstructed were retained for analysis.

Because the infrared window altered thermal transmission and introduced reflections, thermal measurements were calibrated *in situ* using a black reference target measured inside the closed chamber across a known temperature range (approximately 25–45 °C). A linear correction equation derived from these measurements was subsequently applied to all extracted thoracic temperature values.

Thermal analyses were conducted in FLIR Research Studio using an emissivity setting of 0.97 for *B. terrestris* thoracic cuticle, following measurements obtained using the protocol of (Stupski & Schilder, 2021) and consistent with published emissivity values for bee cuticle (0.96–0.99; (Schmaranzer & Stabentheiner, 1988; Stabentheiner & Schmaranzer, 1987; Stupski & Schilder, 2021). Object distance was set to 0.18 m, reflected and atmospheric temperatures were set to ambient temperature (as measured using the respirometry chamber temperature probe), relative humidity was set to ambient relative humidity (as measured using a digital hygrometer), and both transmission and external optics transmission were set to 1.

For each feeding bout, thoracic temperature was measured manually in FLIR Research Studio using a thorax region of interest (ROI). A separate background ROI was placed on a nearby thermally uniform region of the chamber wall to account for background radiation and optical artefacts. Bouts in which reflection conditions differed substantially between pre-feeding and feeding periods were excluded from analyses. Background-corrected thoracic temperature was calculated as:

$$dT=T_{thorax}-T_{background}$$

For each feeding bout, pre-feeding thoracic temperature was estimated from the mean of five frames immediately preceding feeding onset (proboscis contact of the bee with the sucrose solution), whereas feeding temperature was estimated from five evenly spaced frames sampled across the feeding bout. Feeding duration was defined as the interval between initial proboscis contact with the sucrose solution and withdrawal from the feeder. The feeding-associated thoracic temperature response was then calculated as:

$$\Delta\Delta T=dT_{feeding}-dT_{pre-feeding}$$

*Respirometry data analysis*

Respirometry and thermal recordings were synchronised using a shared event consisting of a voltage pulse recorded by the respirometry system and a simultaneous visual marker visible in the thermal recordings. Thermal frame numbers were converted to respirometry time using the synchronisation frame and the thermal camera acquisition frequency (30 Hz).

CO₂ production (V̇CO₂, in mL h⁻¹) was calculated by integrating the differential CO₂ signal (difference between the test and control chambers) over time using the trapezoidal rule. The integrated CO₂ signal was corrected for flow rate (400 mL/min) and bout duration and scaled to an hourly rate.

For each feeding bout, two time windows were defined. Pre-feeding metabolic rate (MR) baseline was defined as the mean CO₂ signal during the 20 s immediately preceding feeding onset. Feeding MR was calculated from the corresponding feeding interval after accounting for system washout delay. Washout delay was estimated experimentally by injecting pulses of air using a microsyringe into the tubing immediately upstream of the chamber and measuring the elapsed time until detection of the resulting CO₂ peak. The mean delay across three injections was 8.99 s, and this offset was applied to align feeding bouts with respirometry traces. Feeding-associated metabolic response was calculated as:

$$\Delta MR=MR_{feeding}-MR_{baseline}$$

*Specific dynamic action (SDA) quantification*

Specific dynamic action (SDA) was quantified from post-feeding CO₂ traces as the increase in metabolic rate associated with digestion and nutrient assimilation, following established definitions of postprandial metabolic responses (Secor, 2009). Continuous CO₂ recordings (ppm; µmol mol⁻¹) were smoothed using a centred rolling mean over a 5 s window to reduce high-frequency noise while preserving biologically meaningful variation in the signal.

For each feeding bout, baseline pre-feeding metabolic rate was defined as the mean CO₂ concentration during the 20 s immediately preceding feeding onset (based on shifted feeding times). Baseline variability was estimated as a robust standard deviation equivalent using the median absolute deviation (MAD × 1.4826), reducing sensitivity to transient fluctuations. Where MAD was zero or undefined, ordinary standard deviation was used instead. To account for potential drift in baseline levels across the recording, a second baseline (“recovered baseline”) was defined as the mean CO₂ concentration during the 20 s immediately preceding the subsequent feeding bout. For final feeding bouts (F3), where no subsequent feeding occurred, the final 20 s of the recording were used provisionally.

A feeding-associated metabolic response was considered present when the maximum smoothed CO₂ concentration within a bounded post-feeding window exceeded the higher of the pre-feeding and recovered baselines by more than 1.5 times the pooled baseline noise, calculated as the median of the pre-feeding and recovered-baseline noise estimates. Peak detection was restricted to a window extending from feeding onset to 60 s after feeding termination. This window captures responses occurring during feeding or shortly after feeding while limiting inclusion of unrelated fluctuations later in the trace and corresponds closely to the maximum feeding duration observed in the dataset (~65 s). Sensitivity analyses confirmed that varying this window (45–75 s) did not alter qualitative outcomes.

Recovery from the SDA response was defined as the first sustained return of CO₂ levels below the recovered baseline plus 1.5 times its associated noise. This condition was required to be maintained continuously for at least 5 s to avoid misclassification due to short-lived fluctuations. Recovery detection began 3 s after the detected peak to prevent premature assignment of recovery.

SDA responses were classified as full when both a feeding-associated peak and subsequent recovery were detected. Responses with a detectable peak but no recovery prior to the next feeding bout (or end of the recording) were classified as partial, while bouts without a detectable peak were classified as non-responses.

The magnitude of SDA was quantified as the total excess CO₂ production above baseline, integrated from feeding termination until recovery or, where recovery was not observed, until the next feeding bout or end of the trace. Integration was performed using trapezoidal numerical methods, with negative deviations from baseline excluded. To provide recovery-independent measures of early SDA dynamics, fixed-window integrals of excess CO₂ were also calculated over the first 30 s, 60 s, and 120 s following feeding.

CO₂ production was converted to volumetric output using measured flow rates. Excess CO₂ production was then converted to energetic units assuming predominantly carbohydrate metabolism. The energetic equivalent of CO₂ production was taken as 20.97 J mL⁻¹ CO₂, derived from the stoichiometry of sucrose oxidation. This assumes a respiratory quotient close to unity, which is consistent with measurements in bees during active behaviours and floral interactions (Rossi et al., 2026; Rothe & Nachtigall, 1989). Meal energy content was calculated from sucrose concentration and solution density following established conversion methods (Pattrick et al., 2025), allowing SDA to be expressed both as absolute energy expenditure and relative to ingested energy.

To assess the robustness of SDA detection and quantification, sensitivity analyses were conducted by varying response and recovery thresholds, recovery-duration criteria, and peak-detection window length. Across all 81 parameter combinations, the direction and relative magnitude of temperature effects on SDA metrics remained unchanged (Supplementary Results; Table S3).

After data processing, the dataset comprised 137 feeding bouts. SDA responses were detected in 109 bouts, all of which reached full measurable scope and were suitable for duration analyses. Additional filtering was applied for specific response metrics: total SDA energy was analysed for responses with positive integrated scope, early-phase SDA (AUC₆₀) for bouts with positive post-feeding area within the first 60 s, and peak amplitude and time-to-peak for responses with valid positive timing values.

*Statistical modelling details*

Statistical analyses were performed in R (v4.4.3; R Core Team, 2024) using the packages *lme4* (Bates et al., 2015), *lmerTest (Kuznetsova et al., 2017)*, *nlme* (Pinheiro et al., 2019), *glmmTMB* (Brooks et al., 2017), *emmeans* (Lenth, 2024), *DHARMa* (Hartig, 2024), and *performance* (Lüdecke et al., 2021).

Body mass was quantified as pre-feeding body mass and log₁₀-transformed and mean-centred prior to analysis. Feeding duration was log-transformed and mean-centred when included as a covariate.

Model assumptions were evaluated using residual diagnostics, quantile–quantile plots, simulated residuals generated using *DHARMa*, and variance inflation diagnostics. Model performance was summarised using marginal and conditional R² values and intraclass correlation coefficients (ICC) where appropriate.

Significant interactions involving continuous covariates were explored using estimated marginal trends (emtrends) implemented in *emmeans*.

*Body-mass sensitivity analyses for metabolic-rate models*

To evaluate the robustness of body-mass effects detected in the metabolic-rate (MR) analyses, we conducted a series of supplementary sensitivity analyses focused on potential leverage from unequal variance structure, extreme body sizes, and model specification. These analyses were performed for both pre-feeding MR and feeding MR mixed-effects models.

First, because residual variance appeared greater at 25°C than at 35°C, we fitted heteroscedastic linear mixed-effects models using a temperature-specific variance structure (varIdent in nlme). These models retained the same fixed- and random-effects structure as the primary models while allowing residual variance to differ between temperature treatments.

Second, to assess sensitivity to extreme body sizes, we repeated analyses after restricting the dataset to the central 90% of the body-mass distribution (excluding observations below the 5th percentile and above the 95th percentile of pre-trial body mass). We additionally performed a targeted high-mass sensitivity analysis in which observations from the 25°C treatment exceeding the global 90th percentile of centred log₁₀ body mass were excluded, because exploratory diagnostics indicated that the steepest scaling relationships occurred primarily among the largest individuals under cooler conditions.

Third, we evaluated whether among-individual variation in body-mass scaling improved model fit by fitting random-slope models that allowed the effect of centred log₁₀ body mass to vary among individuals. These models produced singular fits, indicating insufficient support for stable among-individual variation in scaling slopes, and were therefore retained only as diagnostic checks.

Finally, we tested for nonlinear body-mass scaling by adding quadratic body-mass terms and their interaction with temperature to the feeding MR model. Quadratic terms were not supported, indicating no evidence for substantial nonlinear scaling across the observed body-mass range.

*Classical scaling analyses using F1 only*

To assess whether the observed body-mass relationships were consistent across alternative scaling approaches, we conducted supplementary analyses restricted to the first feeding bout (F1), thereby avoiding repeated-measures structure associated with later feeding events. Within each temperature treatment, relationships between log₁₀-transformed metabolic rate and log₁₀-transformed body mass were analysed using both ordinary least-squares (OLS) regression and standardized major axis (SMA) regression implemented in the smatr package. OLS analyses were included to estimate directional predictive relationships, whereas SMA analyses were used as a classical allometric sensitivity check because both variables may contain measurement and biological variation. These analyses were intended as descriptive robustness assessments rather than primary inferential tests of metabolic scaling exponents.

*Mass-specific metabolic-rate analyses*

Mass-specific metabolic rate (MSMR; mL h⁻¹ g⁻¹) was analysed as a secondary mass-standardised response. These analyses were included to describe metabolic patterns after expressing V̇CO₂ per unit body mass, not as parallel primary model-selection analyses. Because body mass is already incorporated into the MSMR denominator, body mass was not included as a predictor in the primary MSMR models.

Separate models were fitted for pre-feeding MSMR, feeding MSMR, and the change in MSMR during feeding. Fixed effects included ambient temperature, feeding bout, feeding duration, and colony identity, with BeeID included as a random intercept. For pre-feeding and feeding MSMR, responses were log₁₀-transformed prior to analysis. Delta MSMR was analysed on the original scale. Residual body-size dependence was assessed using diagnostic models in which centred log₁₀ body mass was added as an additional covariate to each MSMR model.

*Sensitivity analysis of SDA detection parameters*

To assess the robustness of specific dynamic action (SDA) quantification to parameter choice, we conducted a systematic sensitivity analysis varying key detection and recovery parameters.

SDA metrics were recomputed across all combinations of the following parameter values:

- Response detection threshold (k): 1.0, 1.5, 2.0
- Recovery threshold (k): 1.0, 1.5, 2.0
- Minimum sustained recovery duration: 5, 10, 20 s
- Peak detection window (post-feeding): 45, 60, 75 s

This resulted in 81 unique parameter combinations.

All other aspects of the SDA detection pipeline (baseline definition, smoothing, integration, and temperature normalisation) were held constant.

For each parameter combination, the full SDA detection procedure was repeated, including:

- baseline estimation
- response detection
- peak identification within the bounded window
- recovery detection
- calculation of SDA amplitude, time to peak, duration, and integrated SDA scope

Derived energetic metrics (e.g. SDA energy) were recalculated from excess CO₂ production using the same conversion as in the main analysis.

For each parameter combination, we summarised:

- mean and median SDA energy
- proportion of events with detected responses
- proportion of events reaching recovery

These metrics were calculated separately for each temperature treatment (25°C and 35°C).

Robustness was evaluated based on whether:

1. The direction of temperature effects (25°C vs 35°C) remained consistent
2. The relative magnitude of responses was preserved
3. Detection rates remained stable across parameter combinations

Supplementary Results:

*Body-size audit*

To verify that temperature- and colony-level differences in metabolic and thermal traits were not driven by systematic variation in body size, we tested whether pre-trial body mass differed among colonies or between temperature treatments. A linear model fitted to unique individuals showed no significant effect of colony identity (F2,51 = 0.59, p = .559) or temperature treatment (F1,51 = 0.50, p = .482) on log₁₀-transformed body mass. Mean body mass was similar across colonies (Colony 2: 0.192 ± 0.077 g; Colony 3: 0.195 ± 0.030 g; Colony 4: 0.175 ± 0.042 g) and between temperature treatments (25°C: 0.190 ± 0.051 g; 35°C: 0.183 ± 0.058 g). These results indicate that observed temperature- and colony-associated differences in metabolic and thermal responses were not attributable to systematic differences in body size among groups.

*Pre-feeding thoracic temperature excess*

Pre-feeding thoracic temperature excess was consistently higher at 25°C than at 35°C (Figure S1A; χ²₁ = 43.02, p < .001; estimated contrast = 1.27 ± 0.19°C), ranging from 1.74 ± 1.29°C to 2.03 ± 1.09°C at 25°C, compared with 0.52 ± 0.25°C to 0.70 ± 0.39°C at 35°C.

The relationship between body mass and pre-feeding thoracic temperature excess depended on ambient temperature (Figure S1B; Temperature × Body mass: χ²₁ = 6.74, p = .009). Larger bees exhibited higher pre-feeding thoracic temperature excess at 25°C (3.82 ± 1.29, 95% CI = 1.23–6.40), whereas no association was detected at 35°C (−0.18 ± 0.83, 95% CI = −1.83 to 1.48).

Pre-feeding thoracic temperature excess was also influenced by the interaction between feeding bout and subsequent feeding duration (Figure S1C; χ²₂ = 23.01, p < .001). Bees that subsequently fed for longer exhibited lower pre-feeding thoracic temperature excess during F1 (−0.54 ± 0.13, 95% CI = −0.80 to −0.27) and F2 (−0.52 ± 0.14, 95% CI = −0.80 to −0.23), whereas no relationship was detected during F3 (0.18 ± 0.16, 95% CI = −0.14 to 0.50). Feeding bout also affected pre-feeding thoracic temperature excess overall (χ²₂ = 8.90, p = .012), with lower values during F2 than F3 (post-hoc p = .017). The model explained substantial variation in pre-feeding thoracic temperature excess (marginal R² = .507; conditional R² = .912), with high repeatability among individuals (ICC = .822).

*Pre-feeding metabolic rate*

Pre-feeding metabolic rate (MR) was substantially higher at 25°C than at 35°C across all feeding bouts (Figure S2A; χ²₁ = 44.22, p < .001). Mean pre-feeding MR ranged from 3.77–4.28 mL CO₂ h⁻¹ at 25°C compared with 1.10–1.48 mL CO₂ h⁻¹ at 35°C. The highest pre-feeding MR was observed prior to F3 at 25°C (4.28 ± 3.07 mL h⁻¹), whereas the lowest occurred prior to F3 at 35°C (1.11 ± 0.54 mL h⁻¹).

The relationship between pre-feeding MR, body mass and feeding behaviour was context dependent. Pre-feeding MR increased with body mass overall (χ²₁ = 4.32, p = .038), but this relationship depended on feeding bout and subsequent feeding duration (Figure S2B; Feeding bout × Body mass × Feeding duration: χ²₂ = 6.30, p = .043). Feeding bouts that subsequently lasted longer tended to be preceded by lower pre-feeding MR overall (χ²₁ = 23.71, p < .001), and this negative relationship was stronger at 25°C than at 35°C (Temperature × Feeding duration: χ²₁ = 7.62, p = .006). The relationship between feeding duration and pre-feeding MR also differed among feeding bouts (Figure S2C; Feeding bout × Feeding duration: χ²₂ = 15.15, p < .001), with the steepest negative association during F1 (−0.48 ± 0.07), compared with F2 (−0.34 ± 0.06) and F3 (−0.25 ± 0.07). The model explained substantial variation in pre-feeding MR (marginal R² = .594; conditional R² = .808), with moderate repeatability among individuals (ICC = .527).

*Body-mass sensitivity analyses for metabolic-rate models*

The preceding body-size audit confirmed that body mass did not differ systematically among colonies or temperature treatments, indicating that the primary temperature effects on metabolic and thermal traits were not attributable to group-level differences in size. Nevertheless, because several metabolic-rate models retained interactions involving body mass, additional sensitivity analyses were conducted to assess whether apparent body-mass dependence was robust to residual variance structure and leverage from large individuals. Selected pre-feeding and feeding MR models were re-fitted using heteroscedastic mixed-effects models allowing residual variance to differ between temperature treatments. In both models, residual variance was substantially greater at 25°C than at 35°C, with estimated variance ratios (25°C relative to 35°C) of 1.84 (95% CI: 1.30–2.60) for pre-feeding MR and 1.71 (95% CI: 1.23–2.36) for feeding MR.

Despite this heterogeneity, the principal biological inferences remained unchanged. Pre-feeding MR remained significantly higher at 25°C, and the stronger negative relationship between feeding duration and MR at cooler temperatures was retained. Feeding MR likewise remained substantially higher at 25°C, and the Temperature × Body mass interaction continued to indicate steeper apparent body-mass dependence under cooler conditions.

To assess the influence of extreme body masses, analyses were repeated after restricting the dataset to the central 90% of body masses (0.112–0.272 g), reducing the dataset from 136 to 123 observations. Under this trimmed dataset, all principal temperature and feeding-duration effects remained qualitatively unchanged for both pre-feeding and feeding MR.

However, the Temperature × Body mass interaction in the feeding MR model was no longer supported after trimming (χ²₁ = 1.54, p = .214), indicating that the steeper apparent scaling relationship at 25°C was partly influenced by individuals in the upper tail of the mass distribution. Estimated body-mass slopes at 25°C were reduced relative to the primary analysis, whereas slopes at 35°C remained comparatively shallow.

Because the strongest apparent mass dependence occurred at 25°C, an additional sensitivity analysis removed observations from the 25°C treatment above the global 90th percentile of centred log₁₀ body mass. This reduced the dataset to 127 observations from 52 individuals.

Under this targeted removal, the Temperature × Body mass interaction in the feeding MR model again became non-significant (χ²₁ = 2.66, p = .103), and the estimated slope at 25°C was further reduced. Nevertheless, the strong main effect of temperature on feeding MR remained unchanged, and the negative relationship between feeding duration and feeding MR persisted.

For pre-feeding MR, the primary selected model did not retain a Temperature × Body mass interaction. Instead, body-mass effects remained embedded within interactions involving feeding bout and feeding duration, and these qualitative relationships were broadly preserved across all sensitivity analyses.

Allowing individual-specific body-mass slopes for feeding MR produced a singular fit, with the estimated variance for the random slope collapsing to zero. This indicated that the available data did not support reliable estimation of among-individual variation in body-mass scaling.

Additional analyses incorporating quadratic body-mass terms provided no evidence for nonlinear scaling relationships. Neither the quadratic body-mass term (χ²₁ = 0.02, p = .880) nor its interaction with temperature (χ²₁ = 0.33, p = .564) improved model fit or explained additional variance.

Together, these analyses indicate that the principal temperature effects on metabolic rate were highly robust to alternative variance structures and body-mass filtering procedures. In contrast, apparent temperature-dependent body-mass scaling relationships, particularly for feeding MR at 25°C, were sensitive to leverage from high-mass individuals and should therefore be interpreted cautiously.

*Classical scaling sensitivity analyses using F1 only*

To evaluate whether the observed body-mass relationships were consistent with classical allometric scaling approaches, we additionally analysed first feeding bouts (F1) separately using ordinary least-squares (OLS) and standardized major axis (SMA) regressions within each temperature treatment.

Both approaches recovered positive associations between body mass and metabolic rate at both temperatures. However, estimated scaling exponents were consistently steeper at 25°C than at 35°C and varied substantially between analytical methods. For feeding MR, OLS slopes were 2.22 at 25°C and 0.91 at 35°C, whereas SMA slopes were 4.05 and 1.51, respectively. Similar patterns were observed for pre-feeding MR.

The substantial divergence between OLS and SMA estimates, together with broader confidence intervals and elevated residual variance at 25°C, indicates that scaling estimates were sensitive to model specification and variance structure. These analyses therefore support the qualitative conclusion that larger bees exhibited disproportionately higher metabolic rates under cooler conditions, while also indicating that the magnitude of the apparent scaling relationship should be interpreted cautiously.

*Q_10_-corrected metabolic-rate sensitivity analyses*

To account for variation in ambient temperature between trials within each treatment temperature, V̇CO₂ values were temperature-corrected using a Q₁₀ value of 2 (Lighton, 2008) and the mean ambient temperature recorded for each treatment group (25.13 °C and 35.05 °C). Q_10_-corrected metabolic-rate models produced the same qualitative conclusions as the primary MR analyses. For pre-feeding MR, the final selected Q_10_-corrected model retained the same interaction structure as the uncorrected model, including Temperature × Feeding duration (χ²₁ = 7.62, p = .006), Feeding bout × Feeding duration (χ²₂ = 15.14, p < .001), and Feeding bout × Body mass × Feeding duration (χ²₂ = 6.23, p = .044). Ambient temperature remained a strong predictor of Q10-corrected pre-feeding MR (χ²₁ = 44.02, p < .001), with higher values at 25°C than at 35°C. Model fit was nearly identical to the uncorrected analysis (marginal R² = .592; conditional R² = .808; ICC = .529).

For feeding MR, Q10 correction likewise did not alter the main inference. The final selected Q10-corrected feeding MR model retained Temperature × Body mass (χ²₁ = 5.42, p = .020), Feeding bout × Feeding duration (χ²₂ = 22.43, p < .001), and Body mass × Feeding duration (χ²₁ = 6.20, p = .013), matching the structure and interpretation of the uncorrected feeding MR model. Ambient temperature remained a strong predictor of feeding MR (χ²₁ = 62.59, p < .001), and model fit was again nearly unchanged (marginal R² = .618; conditional R² = .798; ICC = .471). These analyses confirm that the primary metabolic-rate results were robust to Q10 correction.

*Mass-specific metabolic rate*

Mass-specific metabolic rate (MSMR) showed the same broad temperature pattern as absolute metabolic rate, with higher values at 25°C than at 35°C across feeding bouts (Figure S3). Descriptively, pre-feeding MSMR ranged from 18.3 ± 15.5 to 21.7 ± 12.8 mL h⁻¹ g⁻¹ at 25°C, compared with 5.80 ± 2.33 to 8.34 ± 4.06 mL h⁻¹ g⁻¹ at 35°C. Feeding MSMR ranged from 20.0 ± 15.2 to 24.0 ± 13.2 mL h⁻¹ g⁻¹ at 25°C, compared with 6.52 ± 2.67 to 8.86 ± 3.90 mL h⁻¹ g⁻¹ at 35°C.

In the pre-feeding MSMR model, MSMR was significantly higher at 25°C than at 35°C (Figure S3A; χ²₁ = 33.22, p < .001; estimated contrast = 0.421 ± 0.073, p < .001). Feeding bout did not significantly affect pre-feeding MSMR (χ²₂ = 1.74, p = .420), whereas feeding duration was negatively associated with pre-feeding MSMR (χ²₁ = 24.99, p < .001). The model explained substantial variation in pre-feeding MSMR (marginal R² = .349; conditional R² = .731), with moderate repeatability among individuals (ICC = .587). Adding centred log₁₀ body mass as a residual covariate did not reveal significant residual body-size dependence (χ²₁ = 1.84, p = .176).

Feeding MSMR was also significantly higher at 25°C than at 35°C (Figure S3B; χ²₁ = 45.30, p < .001; estimated contrast = 0.435 ± 0.065, p < .001). Feeding bout did not significantly affect feeding MSMR (χ²₂ = 0.60, p = .743), while feeding duration was negatively associated with feeding MSMR (χ²₁ = 16.98, p < .001). The model explained substantial variation in feeding MSMR (marginal R² = .384; conditional R² = .717), with moderate repeatability among individuals (ICC = .540). Residual body-mass dependence was weak and not statistically significant after mass-standardisation (χ²₁ = 3.04, p = .081).

The increase in MSMR during feeding relative to pre-feeding was smaller and more variable (Figure S3C). Delta MSMR tended to be higher at 25°C than at 35°C, but this effect was marginal in the primary model (χ²₁ = 3.65, p = .056; estimated contrast = 0.759 ± 0.399, p = .063). Delta MSMR was not significantly affected by feeding bout (χ²₂ = 4.03, p = .133), but was positively associated with feeding duration (χ²₁ = 5.72, p = .017). Model fit was weaker than for pre-feeding or feeding MSMR (marginal R² = .162; conditional R² = .217), and repeatability among individuals was low (ICC = .066). Adding body mass as a residual covariate did not reveal significant residual body-size dependence (χ²₁ = 1.18, p = .278).

*Results of SDA sensitivity analysis*

Across all 81 parameter combinations tested, results were qualitatively identical:

- SDA energy (both mean and median) was consistently higher at 25°C than at 35°C
- The proportion of events with detected responses was consistently higher at 25°C
- The proportion of events reaching recovery was consistently higher at 25°C

No parameter combination resulted in a reversal of temperature effects for any metric.

These results indicate that SDA detection and derived metrics were invariant to parameter choice within the tested range, and that conclusions regarding temperature effects are not sensitive to reasonable variation in detection thresholds, recovery criteria, or peak window duration.

Supplementary Figures:

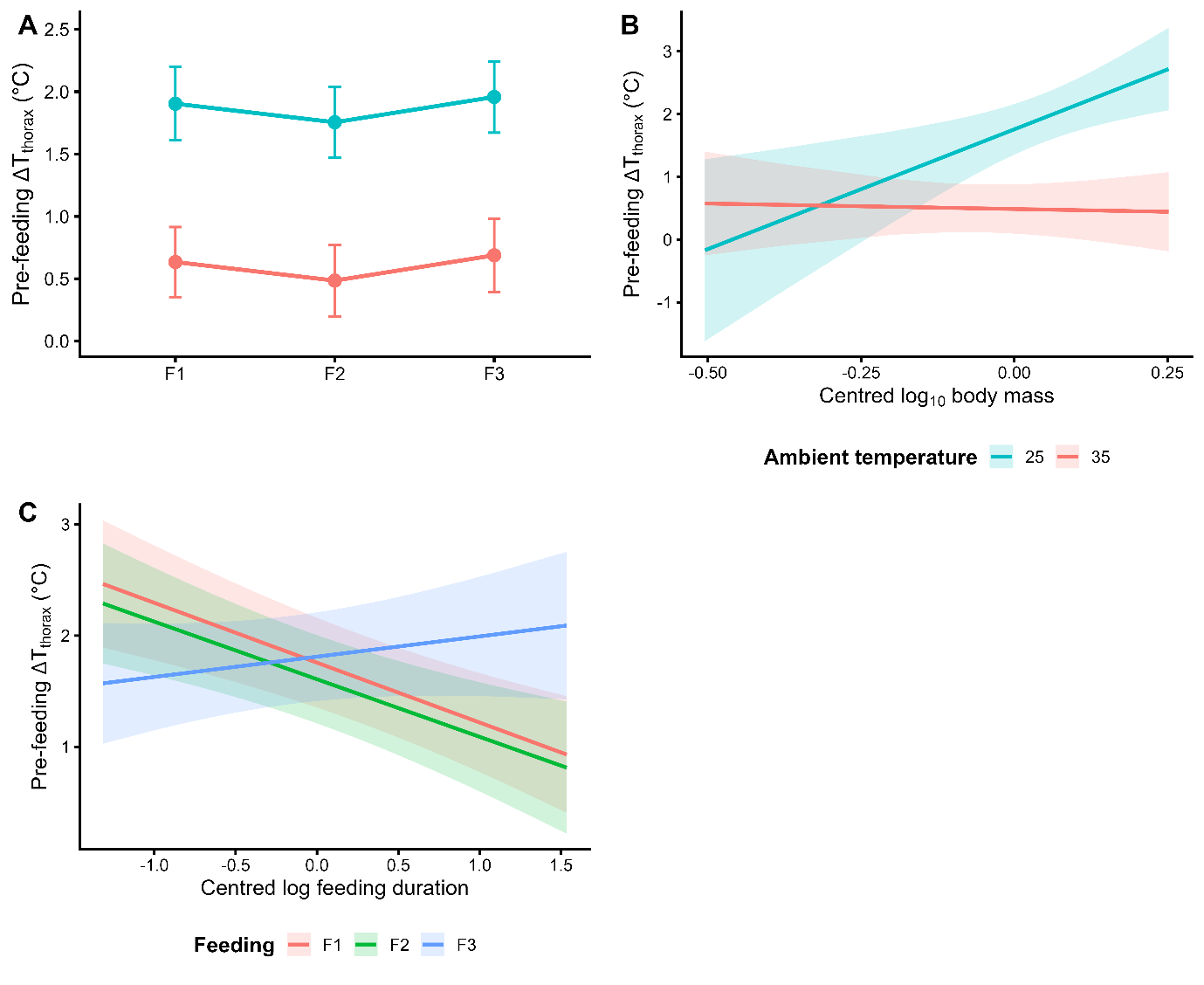

**Figure S1. Pre-feeding thoracic temperature responses and pre-feeding interaction effects across ambient temperatures.** (A) Pre-feeding thoracic temperature excess (ΔT_thorax; thorax − background, °C) across feeding bouts (F1–F3) at 25°C and 35°C. Points represent estimated marginal means ± 95% confidence intervals from the final mixed-effects model. Pre-feeding thoracic temperature excess was consistently higher at 25°C than at 35°C across feeding bouts. (B) Predicted relationship between pre-feeding thoracic temperature excess and centred log₁₀ body mass across ambient temperatures. Lines represent model predictions and shaded ribbons represent 95% confidence intervals from the final mixed-effects model. Larger bees exhibited steeper increases in thoracic temperature excess at 25°C, whereas body-mass effects were weak at 35°C. (C) Predicted relationship between pre-feeding thoracic temperature excess and centred log feeding duration across feeding bouts (F1–F3). Lines represent model predictions and shaded ribbons represent 95% confidence intervals from the final mixed-effects model. Longer feeding events were associated with lower pre-feeding thoracic temperature excess during F1 and F2, whereas no clear relationship was detected during F3. Mixed-effects models included ambient temperature, feeding bout, body mass, feeding duration, and colony as fixed effects, with bee identity included as a random effect. Colours indicate ambient temperature in panels A–B and feeding bout in panel C.

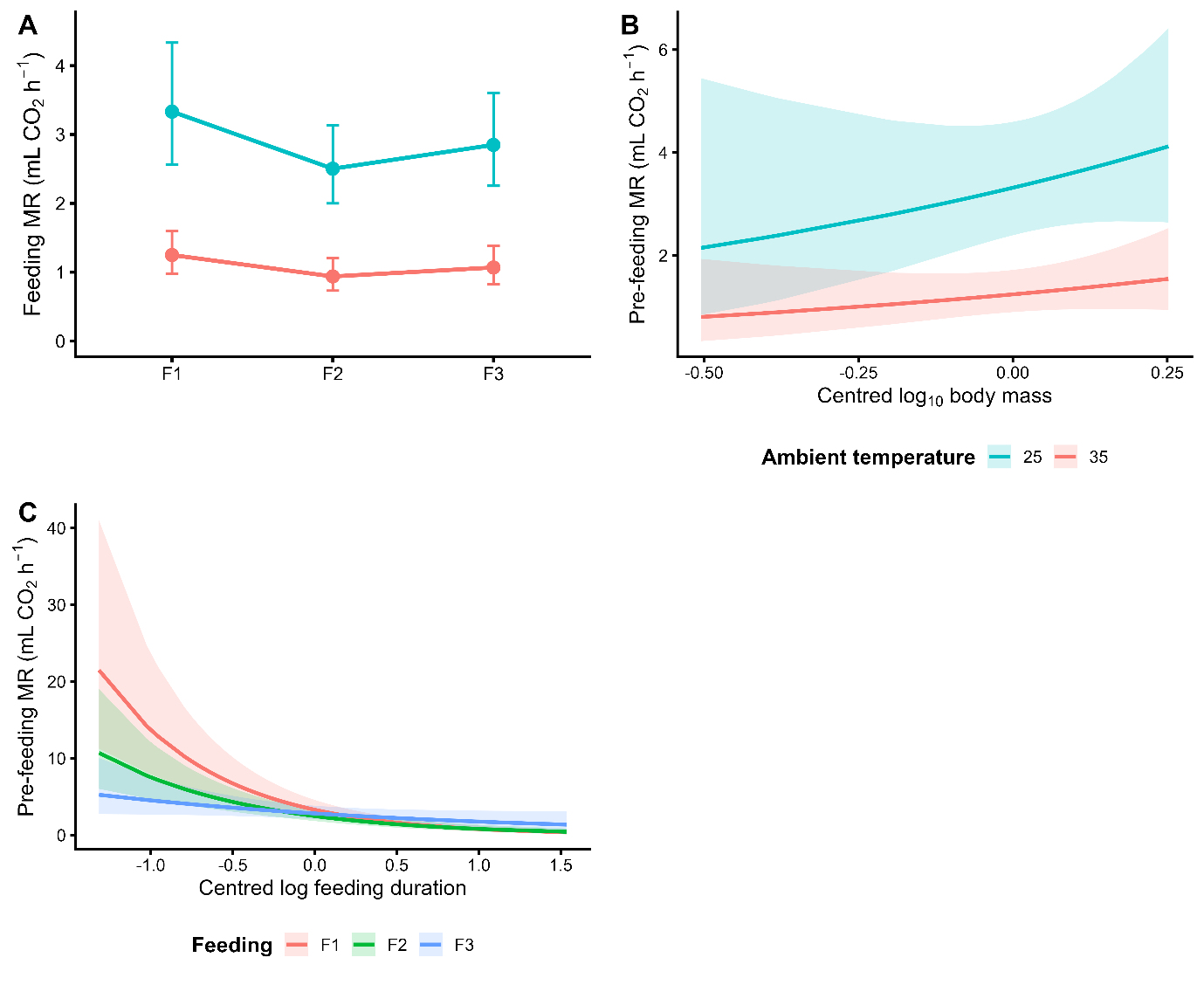

**Figure S2. Pre-feeding metabolic-rate responses and pre-feeding interaction effects across ambient temperatures.** (A) Pre-feeding metabolic rate (MR; mL CO₂ h⁻¹) across feeding bouts (F1–F3) at 25°C and 35°C. Points represent estimated marginal means ± 95% confidence intervals from the final mixed-effects model. Pre-feeding MR was consistently higher at 25°C than at 35°C across feeding bouts. (B) Predicted relationship between pre-feeding MR and centred log₁₀ body mass across ambient temperatures. Lines represent model predictions and shaded ribbons represent 95% confidence intervals from the final mixed-effects model. Larger bees tended to exhibit higher pre-feeding MR, particularly at 25°C, although body-mass effects were weaker and embedded within higher-order interactions involving feeding duration and feeding bout. (C) Predicted relationship between pre-feeding MR and centred log feeding duration across feeding bouts (F1–F3). Lines represent model predictions and shaded ribbons represent 95% confidence intervals from the final mixed-effects model. Longer feeding events were associated with lower pre-feeding MR, with the strongest negative relationship observed during F1 and weaker relationships during later feeding bouts. Mixed-effects models included ambient temperature, feeding bout, body mass, feeding duration, and colony as fixed effects, with bee identity included as a random effect. Colours indicate ambient temperature in panels A–B and feeding bout in panel C.

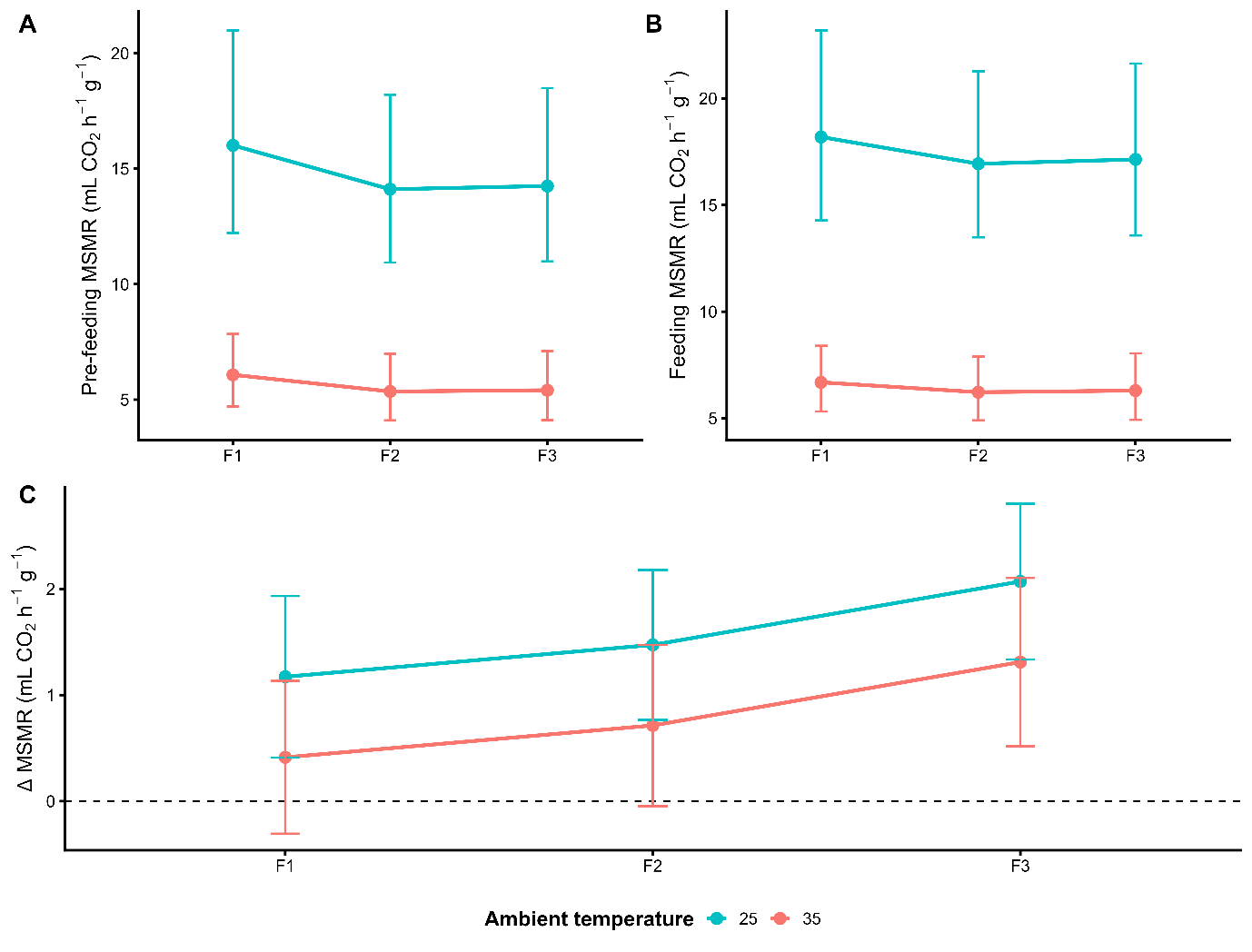

**Figure S3. Mass-specific metabolic rate (MSMR) responses across feeding bouts and temperature treatments.** (A) Pre-feeding MSMR, (B) feeding MSMR, and (C) change in MSMR during feeding relative to pre-feeding (ΔMSMR) across the three feeding bouts (F1–F3) at 25°C and 35°C. Points represent estimated marginal means and error bars indicate 95% confidence intervals from mixed-effects models. MSMR was consistently higher at 25°C than at 35°C for both pre-feeding and feeding metabolic rates, whereas ΔMSMR showed greater variability and weaker temperature dependence.

Supplementary Tables:

**Table S1. Final selected mixed-effects model structures for all analyses**

| **Response variable** | **Final model structure** |
| --- | --- |
| Feeding duration | log_duration ~ Temperature_C * Feeding_bout * log10_mass_c + Colony + (1 \| BeeID) |
| Pre-feeding metabolic rate | log10_baseline ~ Temperature_C + Feeding_bout + log10_mass_c + log_duration_c + Colony + Feeding_bout :log10_mass_c + Temperature_C:log_duration_c + Feeding_bout :log_duration_c + log10_mass_c:log_duration_c + Feeding_bout :log10_mass_c:log_duration_c + (1 \| BeeID) |
| Feeding metabolic rate | log10_feeding ~ Temperature_C + Feeding_bout + log10_mass_c + log_duration_c + Colony + Temperature_C:log10_mass_c + Feeding_bout :log_duration_c + log10_mass_c:log_duration_c + (1 \| BeeID) |
| Change in metabolic rate  during feeding (ΔMR) | delta_mL_h ~ Temperature_C + Feeding_bout + log10_mass_c + log_duration_c + Colony + (1 \| BeeID) |
| Pre-feeding thoracic temperature excess | dT_pre_corr ~ Temperature_C + Feeding_bout + log10_mass_c + log_duration_c + Colony + Temperature_C:log10_mass_c + Feeding_bout :log_duration_c + (1 \| BeeID) |
| Thoracic temperature excess during feeding | dT_feed_corr ~ Temperature_C + Feeding_bout + log10_mass_c + log_duration_c + Colony + Temperature_C:log10_mass_c + Feeding_bout :log_duration_c + (1 \| BeeID) |
| Change in thoracic temperature excess during feeding (ΔTth) | dT_delta ~ Temperature_C + Feeding_bout + log10_mass_c + log_duration_c + Colony + Temperature_C:log10_mass_c + Feeding_bout :log10_mass_c + Temperature_C:log_duration_c + Feeding_bout :log_duration_c + log10_mass_c:log_duration_c + Temperature_C:log10_mass_c:log_duration_c + Feeding_bout :log10_mass_c:log_duration_c + (1 \| BeeID) |
| Change in mass-specific metabolic rate during feeding (ΔMSMR) | delta_MSMR_mL_h_g ~ Temperature_C + Feeding + log_duration_c + Colony + (1 \| BeeID) |
| SDA response occurrence | response_detected ~ Temperature_C + Feeding_bout + log10_mass_c + log_duration_c + Colony + (1 \| BeeID) |
| Full SDA response probability | sda_scope_is_full ~ Temperature_C + Feeding_bout + log10_mass_c + log_duration_c + Colony + (1 \| BeeID) |
| Total SDA energy (strict SDA scope) | sda_energy_j ~ Temperature_C + Feeding_bout + log10_mass_c + log_duration_c + Colony + (1 \| BeeID) + Temperature_C:Feeding_bout + Temperature_C:log10_mass_c + Feeding_bout:log10_mass_c + Temperature_C:log_duration_c + Feeding_bout:log_duration_c + log10_mass_c:log_duration_c + Temperature_C:Feeding_bout:log10_mass_c + Temperature_C:Feeding_bout:log_duration_c |
| SDA early-response magnitude (AUC₆₀) | sda_auc_0_60s_energy_j ~ Temperature_C + Feeding_bout + log10_mass_c + log_duration_c + Colony + (1 \| BeeID) + Temperature_C:Feeding_bout + Temperature_C:log10_mass_c + Feeding_bout:log10_mass_c + Temperature_C:log_duration_c + Feeding_bout:log_duration_c + log10_mass_c:log_duration_c + Temperature_C:Feeding_bout:log10_mass_c |
| Peak SDA amplitude | sda_amplitude_co2 ~ Temperature_C + Feeding_bout + log10_mass_c + log_duration_c + Colony + (1 \| BeeID) + Temperature_C:Feeding_bout + Temperature_C:log10_mass_c + Feeding_bout:log10_mass_c + Temperature_C:log_duration_c + Feeding_bout:log_duration_c + log10_mass_c:log_duration_c + Temperature_C:log10_mass_c:log_duration_c + Feeding_bout:log10_mass_c:log_duration_c |
| SDA duration | sda_duration_s ~ Temperature_C + Feeding_bout + log10_mass_c + log_duration_c + Colony + (1 \| BeeID) |
| Time to peak SDA response | log_time_to_peak_plus1 ~ Temperature_C + Feeding_bout + log10_mass_c + log_duration_c + Colony + (1 \| BeeID) + Temperature_C:Feeding_bout + Temperature_C:log10_mass_c + Feeding_bout:log10_mass_c + Temperature_C:log_duration_c + Feeding_bout:log_duration_c + log10_mass_c:log_duration_c + Feeding_bout:log10_mass_c:log_duration_c |

All models included Bee identity (BeeID) as a random intercept to account for repeated measurements across feeding bouts. Ambient temperature and feeding bout were treated as categorical predictors. Body mass (log10_mass_c) and feeding duration (log_duration_c) were log-transformed and mean-centred prior to analysis.

**Table S2. Summary of body-mass sensitivity analyses for feeding metabolic-rate models**

| **Analysis** | **Temperature effect** | **Body-mass effect** | **Temperature × Body mass** | **Interpretation** |
| --- | --- | --- | --- | --- |
| Primary feeding MR model | Significant | Significant | Significant | Steeper positive mass scaling at 25°C |
| Heteroscedastic variance model | Retained | Retained | Retained | Qualitative inference unchanged after accounting for unequal variance |
| Central 90% body-mass subset | Retained | Retained | Non-significant | Temperature-dependent scaling weakened after excluding mass extremes |
| Removal of largest 25°C individuals | Retained | Retained | Non-significant | Interaction sensitive to high-mass individuals at 25°C |
| Random-slope audit | Retained | Retained | Retained | No support for among-individual variation in scaling slopes (singular fit) |
| Quadratic body-mass audit | Retained | Retained | Retained | No evidence for nonlinear mass scaling |

Across all sensitivity analyses, the dominant effect of temperature on metabolic rate remained robust. Evidence for temperature-dependent body-mass scaling in feeding MR was less stable, however, becoming non-significant after excluding the upper tail of the body-mass distribution. Together, these analyses indicate that although larger bees tended to exhibit higher feeding MR at 25°C, the apparent steepening of scaling relationships under cooler conditions was partly influenced by a small number of high-mass individuals rather than reflecting a uniformly strong population-wide scaling pattern.

**Table S3. Robustness of SDA metrics to parameter choice**

| **Metric** | **Proportion of parameter sets where 25°C > 35°C** |
| --- | --- |
| Mean SDA energy | 1.00 |
| Median SDA energy | 1.00 |
| Response rate | 1.00 |
| Recovery rate | 1.00 |

Across all 81 combinations of response threshold, recovery threshold, sustained recovery duration, and peak detection window, the direction of temperature effects was consistent for all metrics, with SDA responses always higher at 25°C than at 35°C.
